# GB_SynP: a modular dCas9-regulated synthetic promoter collection for fine-tuned recombinant gene expression in plants

**DOI:** 10.1101/2022.04.28.489949

**Authors:** Elena Moreno-Giménez, Sara Selma, Camilo Calvache, Diego Orzáez

## Abstract

Programable transcriptional factors based on the CRISPR architecture are becoming commonly used in plants for endogenous gene regulation. In plants, a potent CRISPR tool for gene induction is the so-called dCasEV2.1 activation system, which has shown remarkable genome-wide specificity combined with a strong activation capacity. To explore the ability of dCasEV2.1 to act as a transactivator for orthogonal synthetic promoters, a collection of DNA parts was created (GB_SynP) for combinatorial synthetic promoter building. The collection includes (i) minimal promoter parts with the TATA box and 5’UTR regions, (ii) proximal parts containing single or multiple copies of the target sequence for the gRNA, thus functioning as regulatory cis boxes, and (iii) sequence-randomized distal parts that ensure the adequate length of the resulting promoter. A total of 35 promoters were assembled using the GB_SynP collection, showing in all cases minimal background and predictable activation levels depending on the proximal parts used. GB_SynP was also employed in a combinatorial expression analysis of an auto-luminescence pathway in *Nicotiana benthamiana*, showing the value of this tool in extracting important biological information such as the determination of the limiting steps in an enzymatic pathway.

## INTRODUCTION

Plant synthetic biology is evolving fast, as high-throughput omics tools provide us with high-quality and precise knowledge about gene expression networks, providing clues for successful engineering interventions. However, there is a shortage of tools capable of controlling the expression of genes in the same precise way as occurs in nature. Many studies still rely on conventional genetic manipulation strategies such as gene knock-out or overexpression driven by constitutive promoters like the *Cauliflower mosaic virus* (CaMV*) 35S* promoter, which could easily cause pleiotropic or even detrimental effects in the transformed organism due to interferences with essential process during their development. To reach its full potential, plant genetic engineering is thus in need of tools for orthogonal and fine-tuned expression of genes. Synthetic promoters are strong allies, not only as tools for gene regulation but also for designing tailor-made metabolic pathways by controlling multiple genes simultaneously.

Plant synthetic promoters typically comprise a minimal promoter and a 5’ regulatory region where cis-regulatory elements are inserted. Regulatory DNA elements are often recruited from the binding sites of natural transcription factors (TFs). The dual architecture of many TFs allows the generation of synthetic TFs that combine their DNA-binding domains with the transcriptional regulatory domains of a different TF and vice versa, creating multiple functional combinations. Moreover, the availability of modular and interchangeable DNA parts greatly expands the possibilities of promoter design. In this regard, modular cloning methods such as MoClo ^1,2^, GoldenBraid ^3^, Mobius Assembly ^4^ or Loop ^5^ facilitate combinatorial rearrangement of promoter elements. GoldenBraid (GB) was conceived as an easy and modular assembly platform based on type IIS restriction enzymes, which makes use of the Phytobricks common syntax ^6,7^ to facilitate the exchangeability of parts. The GB system also proposed a standard measurement using Luciferase/Renilla transient assay to estimate relative expression levels of promoter elements ^8^.

A limitation of this classical approach lies in the hardwired DNA binding specificities of natural TFs, which imposes cis-regulatory elements in a fixed DNA sequence, thus precluding free design, reducing combinatorial power, and comprising full orthogonality. These limitations could be overcome by employing programable transcriptional factors based on CRISPR/Cas9 architecture. The so-called CRISPR activation (CRISPRa) tools, based on the nuclease-deactivated Cas9 protein (dCas9), are becoming commonly used in plants for endogenous gene regulation ^9–12^. The main advantage of CRISPRa tools lays in its programable DNA binding activity, which is encoded in its custom-designed 20-nucleotides guide RNA (gRNA). Another remarkable feature of these tools is their multiplexing capacity, which enables several gRNAs to be directed to the same target gene to ensure higher activation levels, or to target different genes simultaneously to obtain a cascade of activation ^11–13^. CRISPRa tools reported in plants include different protein-fusion strategies, such as SunTag ^14^ and dCas9-TV ^15^, and strategies that make use of modified gRNA scaffolds to anchor additional activator domains ^16,17^. In this last category falls the recently created dCasEV2.1, which showed a strong activation level for endogenous genes that even surpassed those of their natural activation factors ^18^. Interestingly, the transcriptional activation achieved with dCasEV2.1 in *Nicotiana benthamiana* results in remarkable genome-wide specificity. When the promoter region of the endogenous dihydroflavonol-4-reductase (*NbDFR*) gene were targeted for activation in *N. benthamiana* leaves, transcriptomic analysis showed that only the two *NbDFR* homeologous genes were significantly activated, with negligible changes in the rest of the transcriptome. Similar genome-wide specificity was shown for another dCas9-based activation systems ^19^, pointing towards dCasEV2.1 as the ideal system for creating orthogonal synthetic promoters.

In this work, we decided to explore the ability of dCasEV2.1 to transactivate plant genes as a strategy to build a comprehensive collection of orthogonal synthetic promoters. To this end, we chose the 2Kb DNA 5’regulatory region of tomato *SlDFR* gene promoter (p*SlDFR*) as a “model” promoter, given its remarkable inducibility using dCasEV2.1^18^. The strongest activation of p*SlDFR* occurred when targeted at a 20-nucleotides sequence at position −150 from its transcriptional start site (TSS). Taking the p*SlDFR* structure as a prototype, and randomizing most of its sequence, we created a set of synthetic DNA parts comprising distal, proximal and minimal promoter parts (Figure 1A), which, once assembled, produce full orthogonal promoter regions regulated by dCasEV2.1. The promoters in this so-called GB_SynP collection showed negligible basal expression in the presence of unrelated gRNAs, and a wide range of tuneable transcriptional activities. Furthermore, the GB_SynP approach provides a general strategy to generate a virtually endless number of new promoters using interchangeable parts. Such tool can be used for designing large synthetic regulatory cascades where a number of downstream genes (e.g., a whole metabolic pathway) are controlled at custom expression levels by a single programable TF, avoiding repetitive promoter usage. To demonstrate this, we employed GB_SynP promoters in a combinatorial expression analysis of an auto-luminescence pathway in *N. benthamiana* leaves ^20^, extracting valuable information on the limiting steps of the pathway.

**Figure 1.**
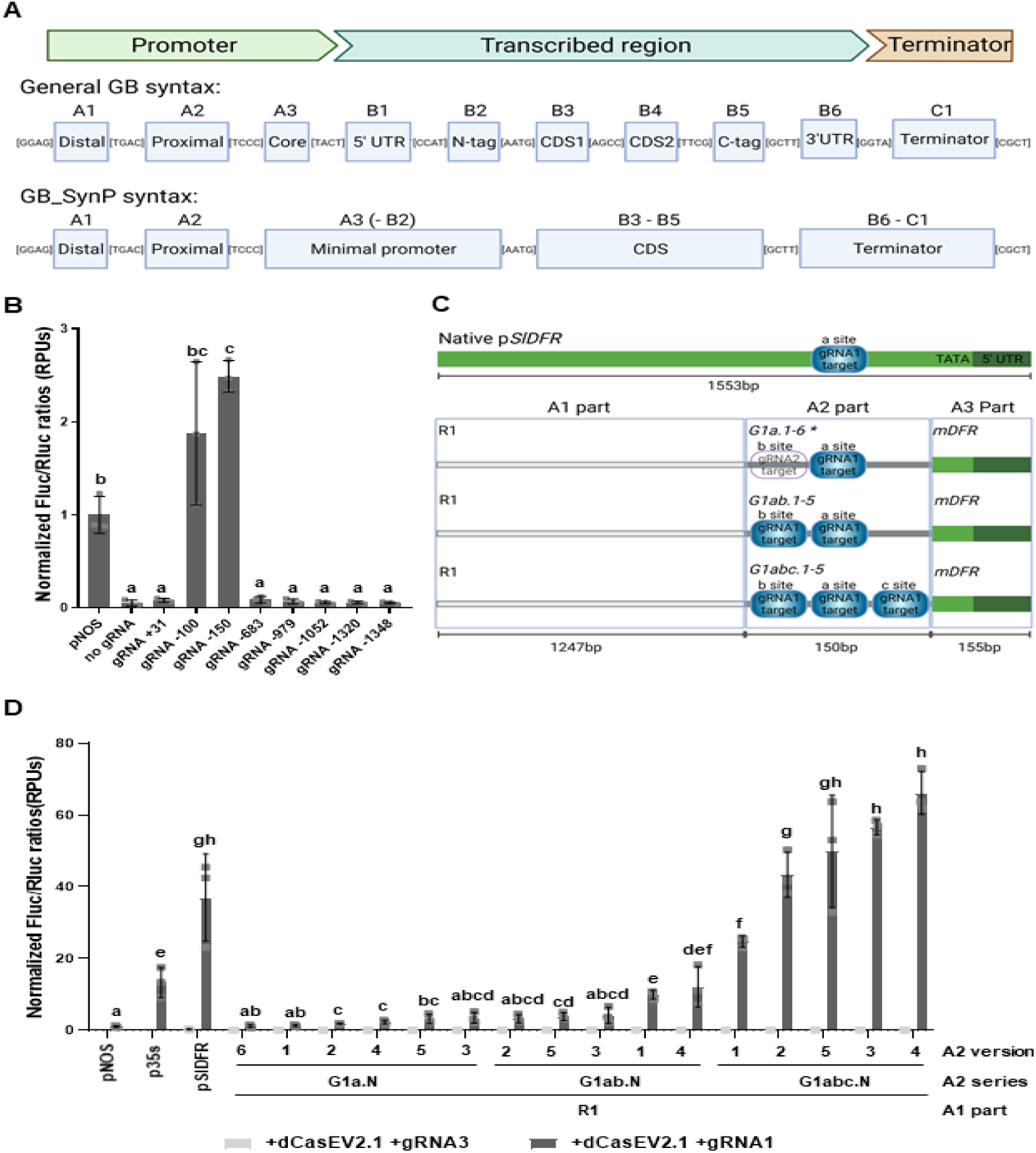
Design and expression range of dCasEV2.1-responsive GB_SynP promoter parts collection. (A) Schematic representation of GoldenBraid (GB) general syntax (A1 to C1 parts), and the specific syntax applied to the GB_SynP collection for A1 distal parts, A2 proximal parts, and A3(−B2) minimal promoter elements. Overhang sequences flanking each part are indicated between brackets. (B) Normalized (FLuc/RLuc) expression levels of *Nicotiana benthamiana* leaves transiently expressing a luciferase reporter gene (FLuc) under *SlDFR* promoter, co-infiltrated with dCasEV2.1 and different gRNAs targeting different positions at the *SlDFR* promoter. Luciferase under *NOS* promoter (p*NOS*) was included as a reference control. (C) Schematic representation of the *SlDFR* promoter (p*SlDFR*) used as a reference for the GB_SynP collection, and the promoter parts designed as A1 distal part (R1), A2 proximal part series containing one (G1a.1-6) two (G1ab.1-5) or three (G1abc.1-5) copies of the target sequence for the gRNA1, and as A3 minimal promoter part (mDFR). Parts in grey indicate the random DNA regions of A1 (light grey) and A2 parts (dark grey). (D) Normalized (FLuc/RLuc) expression levels of *N. benthamiana* leaves transiently expressing FLuc under the regulation of GB_SynP promoters containing R1 and mDFR parts assembled together with the different G1a.N, G1ab.N and G1abc.N A2 parts. Luciferase under *NOS*, CaMV *35s* and Sl*DFR* promoters (p*NOS*, p*35s* and p*SlDFR*, respectively) were included as reference controls. Letters denote statistically significance between (activated) promoters (Student’s t-test, p ≤ 0.05). Error bars represent the average values ± SD (n=3). Figure includes images created with Biorender (biorender.com). *For convenience, the *gRNA2* target was included in position b in A2 parts *G1a*.*1* to *G1a*.*6*, the full name of such parts are *G1aG2b*.*1* to *G1aG2b*.*6*

## RESULTS

### Design of dCasEV2.1-responsive synthetic promoters using the pSlDFR prototype

Previously, we showed in transient transactivation studies *N. benthamiana* that dCasEV2.1 led to a strong transcriptional activation of a Firefly luciferase reporter gene (FLuc) driven by the 2Kb 5’regulatory region of the *SlDFR* promoter (herewith referred to as p*SlDFR*) ^18^. The responsiveness of p*SlDFR* was also confirmed in stably transformed reporter plant lines carrying the pSlDFR:Luc construct, which outperformed other reporter lines employing other promoters. Here, by performing a non-saturated scan of possible target sites in different regions of the p*SlDFR* fragment, we located a 20-nucleotides target box located at position −150 relative to the TSS, named gRNA1, yielding maximum transcriptional activation in transient analysis (Figure 1B). Owing to its proven responsiveness to dCasEV2.1, and especially the low basal expression levels observed in repeated experiments, we decided to use the p*SlDFR* structure as the basis for the design of a new set of dCasEV2.1-regulated synthetic promoters (Figure 1C). A “minimal promoter” element was designed by selecting the region comprising the 5’UTR and the TATA box from the *SlDFR* gene (named mDFR) as previously reported by Garcia-Perez et al. ^21^. This element was assigned a standard A3(−B2) position, according to the Phytobrick syntax, thus being flanked by TCCC and AATG overhangs (Figure 1A). Next to it, several “proximal promoter” parts, assigned to the A2 syntax category, were created. A2 proximal promoters consisted of single or multiple copies of the target sequence for gRNA1 functioning as cis-regulatory boxes, flanked by randomly generated DNA sequences (A2 parts sequences are collected and aligned in Figure S1A). The gRNA1 target in the synthetic parts was maintained at position −150 relative to the TSS (herewith referred as “a site”), mimicking the structure of the native p*SlDFR*. To expand the availability of unique A2 parts and avoid repetitions in promoter choice, six different A2 parts were initially designed (named G1a.1 to G1a.6). These A2 parts contain a single gRNA1 target site at this “a site” (position −150) and each one has a different random background sequence. For convenience, the target sequence of a different gRNA (named gRNA2), was also included in all G1a.N parts at position −210 from the TSS (referred to as “b site”). Later, the collection was further expanded with a second series of five new A2 parts (G1ab.1 to G1ab.5 parts), where a repetition of the gRNA1 box was inserted at the “b site” (position −210). Finally, a third group of five A2 sequences (G1abc.1 to G1abc.5) was created containing the target sequence three times, with the third copy located at position −100 (referred as the “c site”) from the TSS (G1abc.1 to G1abc.5). To finalize the promoter design, an A1 “distal promoter” part (named R1) consisting of 1240 nucleotides of random DNA sequence was designed to mimic the length of the native p*SlDFR*. All randomly designed A1 and A2 sequences were analysed with the TSSP software (http://www.softberry.com/berry) to ensure the absence of spurious cis-regulatory elements.

Promoter parts were next assembled to generate a total of 16 synthetic promoters, which were subsequently combined with the FLuc coding sequence and the CaMV *35s* terminator. All the resulting transcriptional units were further combined with Renilla luciferase (RLuc) under CaMV *35s* promoter for normalization (as required for the standard Luciferase/Renilla transient assay) and the *P19* silencing suppressor and subsequently assayed in transient transactivation experiments in *N. benthamiana* leaves. All promoters showed negligible basal expression levels when co-transformed with a dCasEV2.1 loaded with a gRNA (named gRNA3) which target sequence is not present in the sequences of the promoters. Interestingly, basal expression levels were even lower than those observed with the native p*SlDFR* promoter in all cases tested, probably reflecting the enhanced orthogonality of random A1 and A2 regions. On the contrary, co-transformation with gRNA1 led to substantial transcriptional activation in all promoters assayed, yielding a range of activation levels that increased with the number of copies for the gRNA target present in the A2 element (Figure 1C). Promoters that included the target sequence for gRNA1 once (G1a.N series) showed luciferase levels similar to those obtained with a *NOS* promoter used for normalization and set at a value of 1.0 relative promoter units (RPUs) ^8,13^. Promoters with the target sequence present three times (G1abc.N series) reached activation levels similar to the native p*SlDFR* promoter with values around 50 RPUs on average. The G1ab.N promoter series showed intermediate transcription levels, similar to those obtained with *CaMV* 35s promoter, when activated with dCasEV2.1.

### Expanding the combinatorial GB_SynP collection with additional configurations of the synthetic cis-regulatory region

The proposed modular GB_SynP structure allows, in principle, a limitless extension of the gRNA1-responsive promoter collection by the addition of new distal (using A1 syntax) and minimal promoter (with A3-B2 syntax) parts. To test this, two new A1 distal elements (R2 and R3) with random DNA sequences different to R1 were designed. These new parts were assayed in combination with A2 proximal promoters described above having one (G1a.1), two (G1ab.1) or three (G1abc.1) repetitions of the target sequence for gRNA1 (Figure 2A). As observed in Figure 2B, random distal promoter sequences had no significant influence on the transcriptional levels obtained with GB_SynP promoters. For all promoters assayed, the only relevant factor strongly determining the luciferase activity was the number of cis gRNA1 elements present in the proximal promoter region, proving the orthogonality of distal promoter parts in the GB_SynP design.

**Figure 2.**
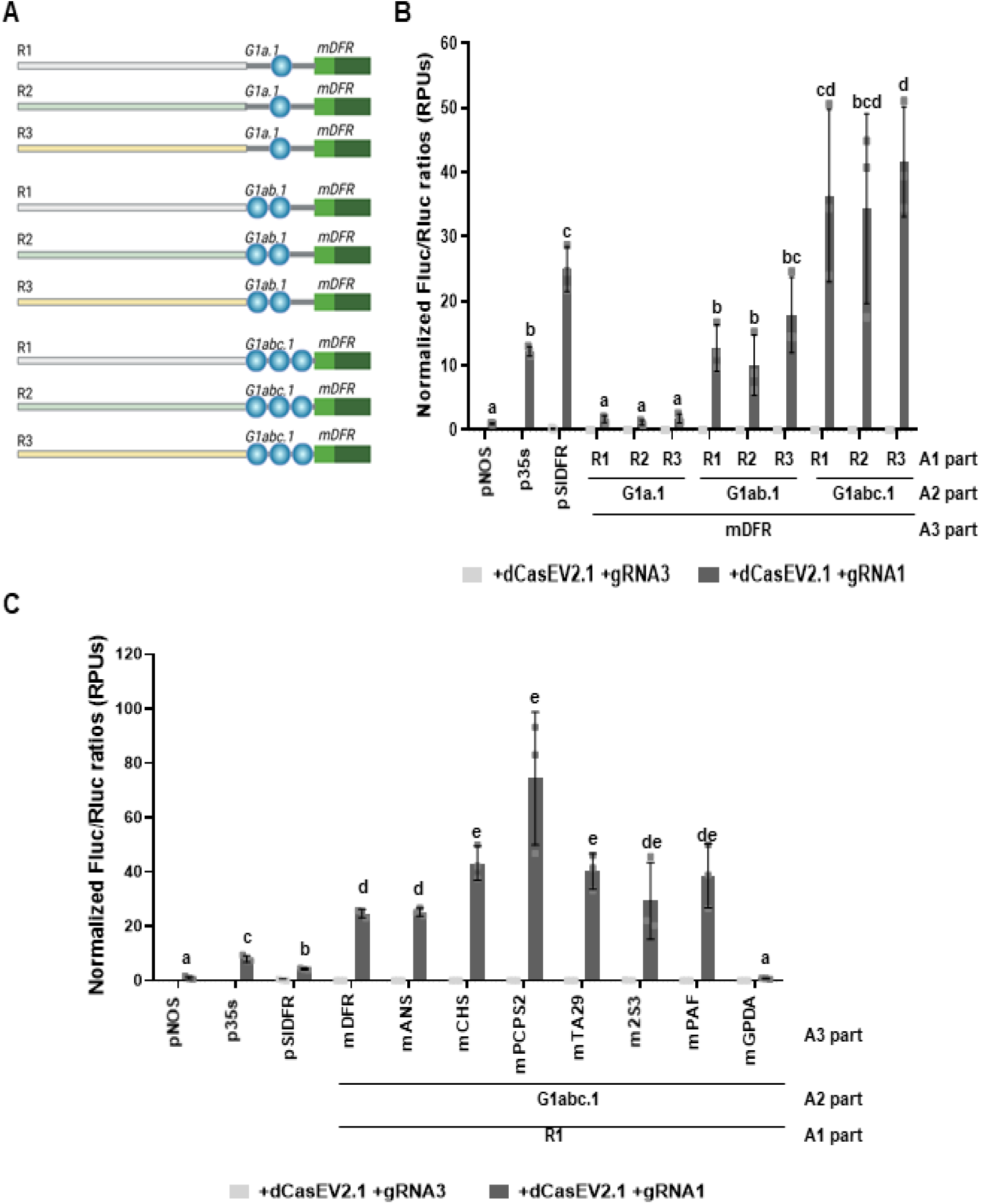
Addition and testing of new A1 distal sequences and A3 minimal promoter parts for the GB_SynP collection. (A) Schematic representation of the GB_SynP promoter series assembled to test the A1 distal parts R1, R2 and R3. A1 parts were combined with A3 mDFR and three different A2 proximal parts containing one (G1a.1) two (G1ab.1) or three (G1abc.1) copies of the gRNA1 target sequence (blue dots). (B) Normalized (FLuc/RLuc) expression levels of *Nicotiana benthamiana* leaves transiently expressing a luciferase reporter gene (FLuc) under the regulation of GB_SynP promoters combining the different A1 distal parts (R1, R2 or R3) with R1 part and different A2 parts including different repetitions of the gRNA1 target. (C) Normalized (FLuc/RLuc) expression levels of *Nicotiana benthamiana* leaves transiently expressing FLuc under the regulation of different GB_SynP promoters assembled with R1 and G1abc.1 parts in combination with different A3 minimal promoter elements. Luciferase under *NOS*, CaMV *35s* and *DFR* promoters (*pNOS, p35s* and pSlDFR, respectively) were included as reference controls. Letters denote statistically significance between (activated) promoters (Student’s t-test, p ≤ 0.05). Error bars represent the average values ± SD (n=3). Figure includes images created with Biorender (biorender.com).

Next, new A3 minimal promoter parts were also added to the collection and functionally assayed. Minimal promoter elements were designed based on the sequences of different strongly-regulated and/or tissue-specific genes from *Solanum lycopersicum, Nicotiana tabacum* and *Arabidopsis thaliana*. In addition, two minimal promoters based on fungal sequences were also created. Table S1 summarizes the genomic regions selected as A3 parts. All minimal promoters were assembled upstream with R1 and G1abc.1 (3xgRNA1-target) parts, downstream with the Luc/Ren reporter, and tested functionally. As shown in Figure 2C, minimal promoters had a stronger influence than A1 distal parts in determining the final transcriptional activity. We observed significant differences (up to 4-fold on average) among the plant promoters assayed. Maximum activation levels corresponded to the mPCPS2 A3 element. Fungal mGPDA showed almost no activity in *N. benthamiana*, however fungal mPAF A3 part promoted high transcriptional levels, similar to other promoter regions obtained from plants.

Despite the expression differences found employing different minimal promoters in the GB_SynP design, the A2 proximal region carrying the dCasEV2.1 cis protospacer elements concentrates most of the regulatory activity. Therefore, it was interesting to investigate modifications in its structure that could accommodate additional regulatory features. Accordingly, we first analysed the influence of the relative position of the cis gRNA1 target to the TSS. New A2 proximal parts were thus designed which included the target for gRNA1 at positions −120 (named “d site”), −150 (“a site”), −210 (“b site”) and −320 (named “e site”) upstream of the TSS (named G1d.1, G1a.7, G1b.1 and G1e.1, respectively, see Figure 3A). As observed in Figure 3B, the transcriptional levels peaked when the gRNA1 target was at positions −120 and −150 from TSS, without statistical differences between these two positions, while the expression decreased when the target was positioned further away from the TSS. For G1e.1 part, which contained the target at “e site” (−320 from TSS), the activated expression levels were ten times lower than the *NOS* promoter used as reference, reaching values of 0.04 RPUs.

**Figure 3.**
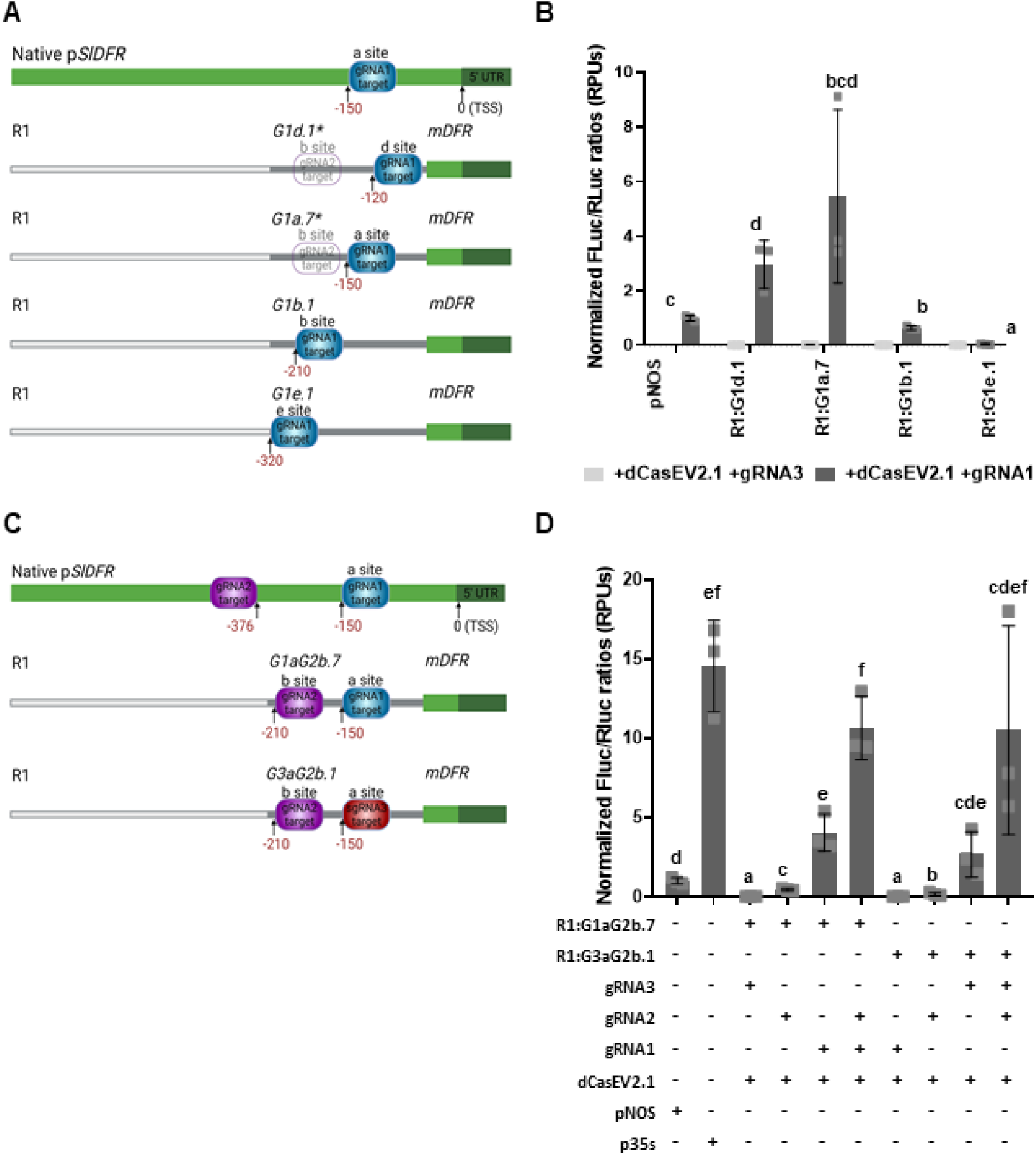
Variation of the cis-regulatory boxes within the A2 proximal parts of the GB_SynP collection. (A) Schematic design of the GB_SynP promoter series containing the target sequence for gRNA1 at position −120 (“d site”), −150 (“a site”), −210 (“b site”) or −320 (“e site”) from the Transcriptional Start Site (TSS). (B) Normalized (FLuc/RLuc) expression levels of *Nicotiana benthamiana* leaves transiently expressing a luciferase reporter gene (FLuc) under the regulation of GB_SynP promoters containing R1 and mDFR parts, in combination with A2 parts including the gRNA1 target sequence in different positions (G1d.1, G1a.7, G1b.1 and G1e.1). Luciferase under *NOS* promoter (p*NOS*) was included as reference. (C) Schematic design of the GB_SynP promoters including the A2 parts G3aG2b.1 and G3aG2b.1. These two A2 parts contain the same sequence except for the gRNA target at position −150 from the TSS (“a site”) which in G1aG2b.7 corresponds to gRNA1 target sequence and G3aG2b.1 part corresponds to gRNA3 target sequence. (D) Normalized (FLuc/RLuc) expression levels of *Nicotiana benthamiana* leaves transiently expressing FLuc under the regulation of GB_SynP promoters assembled with R1 and mDFR parts, in combination with the A2 part G1aG2b.7 or G3aG2b.1. Luciferase under *NOS* and *35S* promoters (p*NOS* and p*35s*, respectively) were included as references. Letters denote statistical significance between signals (Student’s t-test, p ≤ 0.05). Error bars represent the average values ± SD (n=3). Figure includes images created with Biorender (biorender.com). *For convenience, the gRNA2 target was included in position b in A2 parts *G1d*.*1* and *G1a*.*7*, the full name of such parts will be *G1dG2b*.*1* and *G1aG2b*.*7*, respectively.

Next to the position of the target sequence, we analysed the inclusion of new cis-regulatory elements other than gRNA1. For this, we chose the target sequence of gRNA3 as a new cis element, which is natively present at position −161 in the *NOS* promoter. This was previously shown to produce high activation of the *NOS* promoter when targeted with dCasEV2.1 ^18^. We then designed a new proximal element with the exact same sequence as G1a.7 but replacing the gRNA1 target by gRNA3 target (see Figure 3C, A2 parts sequences are collected and aligned in Figure S1B). In both G3a.1 and G1a.7 parts, the target sequence for the gRNA2 at the “b site” (position −210bp) was also present (thus renamed as G1aG2b.7 and G3aG2b.1, respectively). This target sequence is found at position 376 upstream of the TSS in the *SlDFR* promoter and showed low activation in the native promoter ^18^, which could be due to its distance from the TSS. The new A2 parts were then combined with R1 and mDFR parts (Figure 3C), and the resulting full promoters were assayed using single guide or double guide combinations (gRNA1+gRNA2 for G1aG2b.7 promoter, and gRNA3+gRNA2 for G3aG2b.1 promoter). Figure 3D shows that gRNA2 alone triggered a lower response when compared with gRNA1 in G1aG2b.7 or gRNA3 in G3aG2b.1, but still reaching transcriptional values close to a standard *NOS* promoter. gRNA3 and gRNA1 showed similar activation levels when used alone to activate each (4.04 RPUs for gRNA1 in G1aG2b.7 and 2.68 RPUs gRNA3 in G3aG2b.1), while double activation using gRNA2+gRNA1 for G1aG2b.7 and gRNA2+gRNA3 for G3aG2b.1 resulted in higher activation levels (10.63 and 10.05 RPUs, respectively) when compared to using each gRNA individually.

### Combining additional activation domains with dCas9 to activate synthetic promoters

The dCasEV2.1 system is considered to be a second-generation CRISPRa tool ^12^ as it combines the use of two proteins, dCas9 and MS2, to which two activation domains are fused (EDLL and VPR, respectively). This modular architecture can be exploited as an additional source of variability in the system, incorporating different activation domains (e.g., non-viral domains) to the dCas9 and MS2 modules, thus expanding the range of trans-activators for GB_SynP promoters. In addition, other dCas9-based transactivation strategies, such as the SunTag system can be also incorporated. In the SunTag approach, activation domains are fused to a single-chain variable fragment (ScFv) antibody, which in turn binds to a SunTag multiepitope peptide fused to dCas9 protein. To explore these additional expansions of the system, we assayed the two activation domains, ERF2 and EDLL, in four different combinations with dCas9 and MS2 modules, as well as the dCas9:SunTag system with EDLL, ERF2 or VPR fused to the ScFv antibody. All these dCas9-based TFs were co-infiltrated with the reporter R1:G1abc.1:mDFR:FLuc and transiently assayed (Figure 4). Significant activation levels were obtained compared to the background levels in all cases except for those in which ERF2 acted as the main activation domain. The higher activation levels were obtained with the combination dCas:EDLL-MS2:ERF2 and dCas:SunTag-ScFv:VPR, which showed activations of 40-fold and 10-fold respectively, reaching activation levels of 0.75 and 0.14 relative promoter units (RPUs). In all new combinations, the expression levels were similar or lower than the standard p*NOS* signal. The original dCas:EDLL-MS2:VPR (dCasEV2.1) was the only combination that reached expression levels comparable to the CaMV *35s* promoter, thus confirming the unique characteristics of this activation tool in plants.

**Figure 4.**
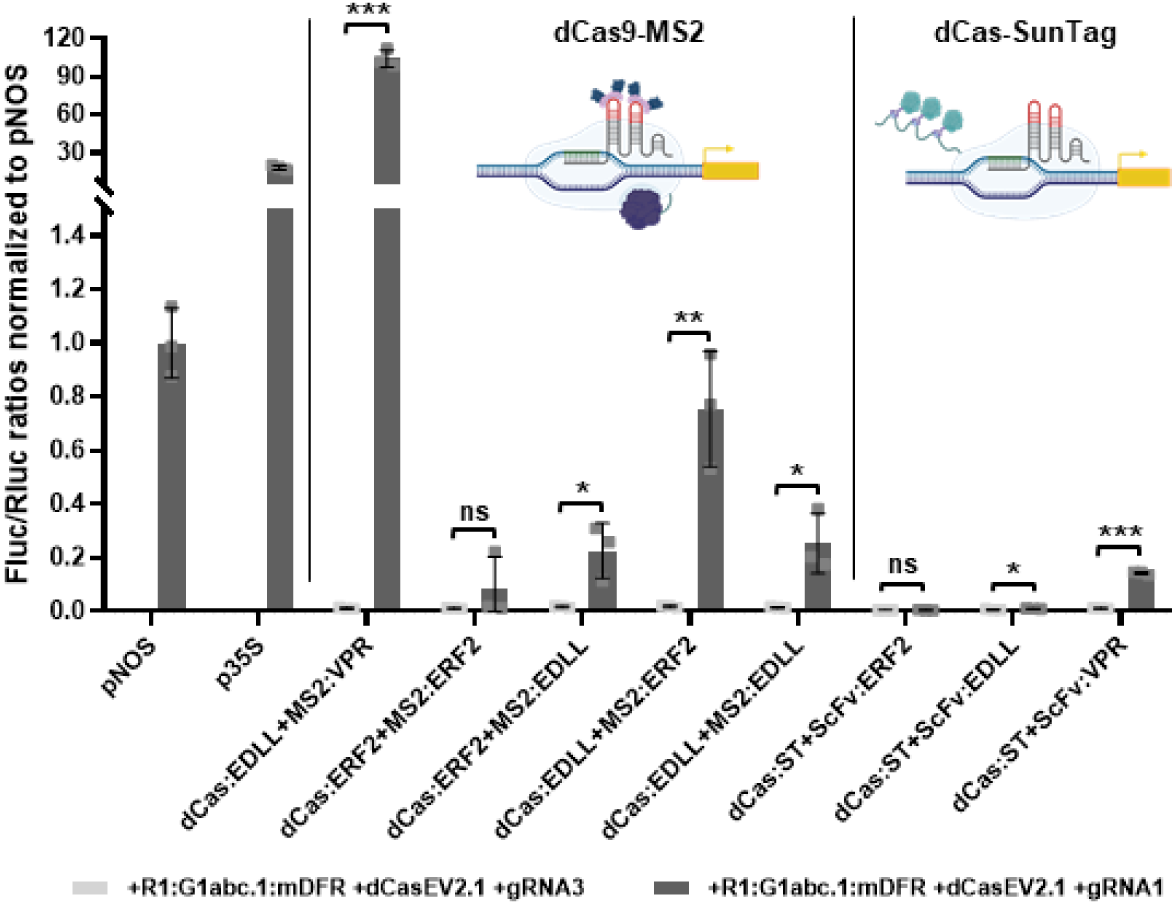
Transactivation of GB_SynP promoters with different CRISPRa strategies. Normalized (FLuc/RLuc) expression levels of *Nicotiana benthamiana* leaves transiently expressing a luciferase reporter gene (FLuc) under the regulation of a GB_SynP promoter containing the A1 part R1, A2 part G1abc.1 and A3 part mDFR (R1:G1abc.1:mDFR), co-infiltrated with dCas9-SunTag or dCas9-MS2 systems harbouring different activation domains. Luciferase under *NOS* and CaMV *35s* promoters (p*NOS* and p*35s*, respectively) were included as references. Asterisks denote statistical significance between activated and basal expression levels, following APA’s standards (Student’s t-test, ns = p < 0.05, * = p ≤ 0.05, ** = p ≤ 0.01 and *** = P ≤ 0.001). Error bars represent the average values ± SD (n=3). Figure includes images created with Biorender (biorender.com).

### Fine-tuning the expression of an auto-luminescence pathway

The fungal auto-luminescence pathway LUZ, described previously by Kotlobay et al. ^22^, was recently adapted to plants ^20,23^. The LUZ pathway has as major advantage that uses the plant’s endogenous caffeic acid as a substrate to produce luciferin, thus avoiding the need for exogenous addition of luciferin substrate. Moreover, the self-sustainable luminescence emission implies that non-destructive assays can be performed, allowing for instance the visualization of time-course kinetics. The pathway comprises four genes, named *HispS* (hispidin synthase), *H3H* (hispidin-3 hydroxylase), *Luz* (luciferase) and *CPH* (caffeylpyruvate hydrolase). *HispS* encodes for the larger enzyme of the pathway which catalyses three consecutive reactions to convert caffeic acid into hispidin, which is then turned into luciferin by a reaction catalysed by *H3H* enzyme. Finally, luciferin is used by *LUZ* enzyme as a substrate to create a high energy intermediate that emits light upon its degradation to caffeylpyruvic acid. The fourth enzyme of the pathway, *CPH*, is included to recycle this degradation product back to caffeic acid, thus closing the cycle (Figure 5A).

**Figure 5.**
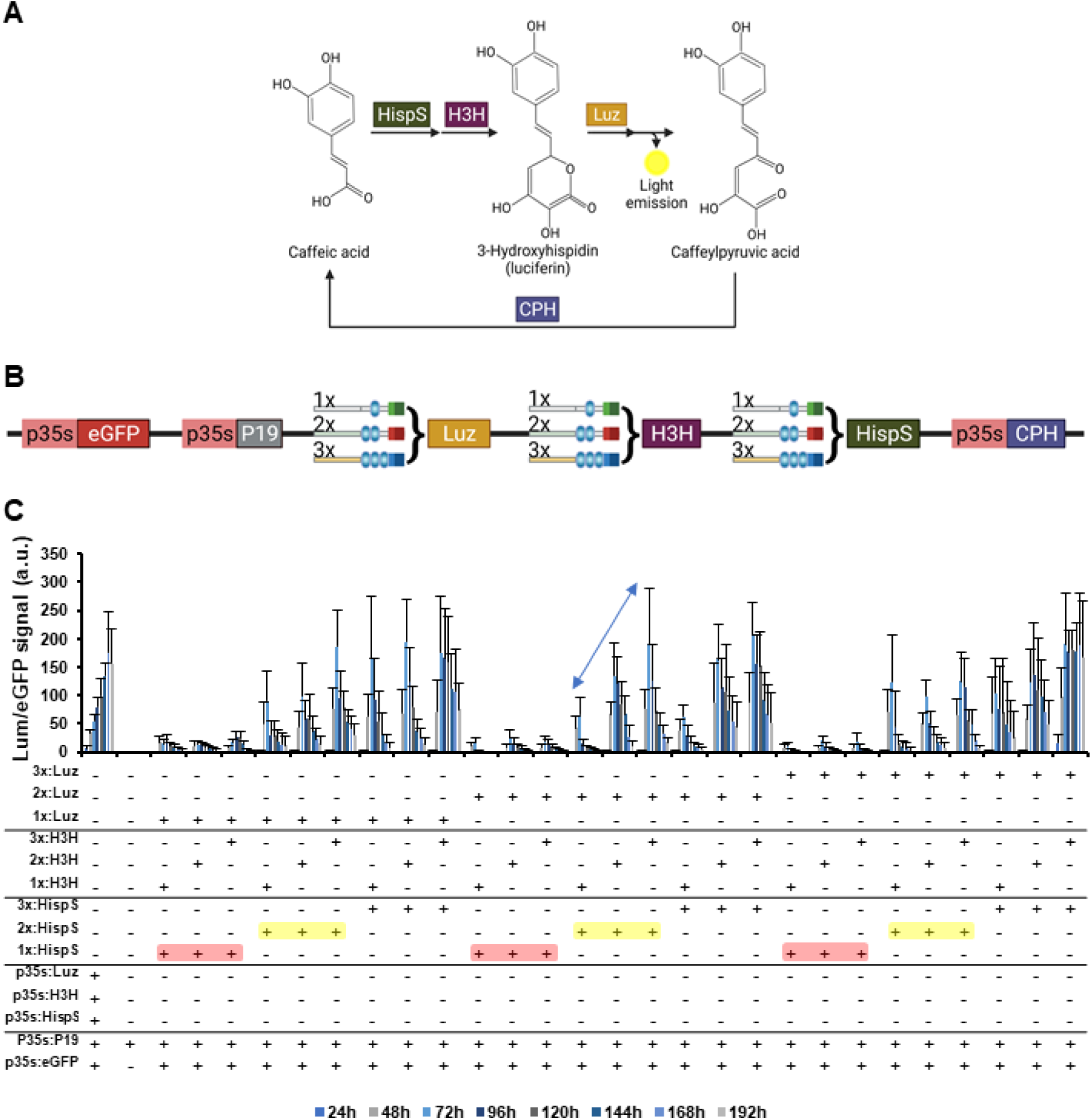
Transient expression of the auto-luminescence LUZ pathway under the regulation of GB_SynP promoters in *Nicotiana benthamiana* leaves. (A) Schematic representation of the LUZ pathway described by Kotlobay et al. ^22^. The pathway consists of three genes (*HispS, H3H* and *Luz*) that converts caffeic acid into caffeylpyruvic acid with the emission of light, and a fourth gene (*CPH*) that turns caffeylpyruvic acid back to caffeic acid. (B) Schematic view of the genetic constructs assembled for expressing the LUZ pathway under the regulation of GB_SynP promoters. *Luz, H3H* and *HispS* genes were assembled in combination with synthetic promoters having one (1x, corresponds to promoter R3:G1a.1:mDFR), two (2x, corresponds to promoter R2:G1ab.1:m2S3) or three (3x, corresponds to promoter R1:G1abc.1:mPCPS2) targets for *gRNA1*. The fourth gene of the pathway, *CPH*, was constitutively expressed in all combinations under a CaMV *35s* promoter (p*35s*). The constructs included a constitutively expressed enhanced GFP protein (p35s:eGFP) for normalization of the luminescence values, and the P19 silencing suppressor (p35s:P19). (C) Time-course expression of the 27 constructs expressing the LUZ pathway transiently in *N. benthamiana* leaves under the regulation of 1x, 2x or 3x GB_SynP promoters, co-infiltrated with dCasEV2.1 system and gRNA1. A constitutive control was included with all four genes of the LUZ pathway expressed under p*35s*. A negative control was also included by infiltration of P19 silencing suppressor (p35s:P19). Luminescence (Lum) values were normalized using fluorescence values produced by the constitutively expressed eGFP (p35s:eGFP) included in all the constructs as an internal control. Red boxes indicate combinations highlighted in the text where *HispS* is under regulation of 1x (gRNA-target) promoter, while yellow boxes indicate combinations where it is expressed under 2x (gRNA-target) promoter. The arrow highlights the three combinations where *HispS* and *Luz* are under regulation of 2x (gRNA-target) promoter, being *H3H* the only gene regulated by a different promoter in each of those three combinations. Error bars represent the average values ± SD (n=12). Figure includes images created with Biorender (biorender.com).

Adapting the LUZ pathway as a reporter for gene expression analysis in plants requires identifying which genes in the pathway act as limiting steps, so that changes in their transcriptional levels are directly translated into changes in light intensity. Therefore, to understand the limiting steps governing the expression of this pathway in *N. benthamiana*, we took advantage of the combinatorial power and the wide expression range of the GB_SynP tool and created a series of assemblies to differentially regulate the expression of the *Luz, H3H* and *HispS* genes. We used three different GB_SynP promoters having either one (R3:G1a.1:mDFR, 1x gRNA-target), two (R2:G1ab.1:m2S3, 2x gRNA-target) or three targets (R1:G1abc.1:mPCPS2, 3x gRNA-target) for the gRNA1 (Figure 5B, see Figure S2 for the strength of each promoter). The *CPH* recycling enzyme was kept under the constitutive CaMV *35s* promoter in all genetic constructs to reduce the complexity of the analysis. An enhanced GFP protein (eGFP) under the CaMV *35s* promoter was also included in each construct to serve as internal reference for normalization. The normalized luminescence values of the resulting 27 pathway combinations co-infiltrated with dCasEV2.1 and gRNA1 are depicted in Figure 5C. The figure shows a time-course from day 1 to day 7 for each synthetic pathway, taking advantage of non-destructive auto-luminescence measurements. As expected, the highest luminescence values, comparable to those obtained when all three enzymes are controlled by the constitutive CaMV *35S* promoter, were reached when all three genes in the pathway were regulated by 3x gRNA-target promoters, while the lowest transcriptional levels were found when the three genes were under 1x gRNA-target promoter. Almost no luminescence was observed for any of the 27 combinations when co-infiltrated with dCasEv2.1 and an irrelevant gRNA (gRNA3, see Figure S3).

The analysis of the remaining pathway combinations served as guidance to understand the regulation of the synthetic pathway. In all combinations where *HispS* was driven by 1x gRNA-target promoters, the resulting normalized luminescence values remained at basal levels regardless of the synthetic promoters used to regulate the remaining genes (see red boxes in Figure 5C), thus indicating that *HispS* expression acts as a limiting factor. Raising *HispS* levels to those provided by dCasEV2.1-activated 2x gRNA-target promoters was sufficient to prove the effects of the regulation of the remaining genes (see yellow boxes in Figure 5C). Particularly informative for reporting applications are those combinations where 2x gRNA-target promoters regulate both *Luz* and *HispS* (see arrow in Figure 5C). Using this conformation, the modifications in the promoter strength driving *H3H* are readily reflected in luminescence levels following a positive lineal trend with no signs of saturation. Considering that activated 2x gRNA-type promoter show expression levels in the range of a *NOS* promoter, this indicates a reporter system with an appropriate dynamic range could consist in a pathway where *Luz* and *HispS* are regulated by constitutive *NOS* promoter and *H3H* is set under a variable-strength promoters for e.g., transactivation studies.

## DISCUSSION

The synthetic promoters whose expression is regulated via CRISPRa systems are promising orthogonal tools for Synthetic Biology. CRISPRa-based synthetic promoters have been previously reported in bacteria ^24^, yeast ^25^ and human cells ^25,26^, being GB_SynP one of the first collections reported in plants, together with the work recently published by Kar et al. ^27^. In contrast to the commonly used activation systems based on transcription activator-like effectors (TALEs) or the zinc finger proteins, which require re-engineering of the DNA-binding motifs for each target sequence ^28,29^, GB_SynP allows the creation of promoters with completely different cis-boxes by simply creating new A2-type cis-regulatory parts, and its corresponding gRNA transcriptional unit, both elements being only a few hundred base pairs long. Here, we also demonstrated that two completely different gRNAs (gRNA1 and gRNA3) can reach similar activation levels when positioned in the same position within the GB_SynP synthetic promoter, implying that potentially any 20-nucleotides sequence can be used as a cis-regulatory box. Specificity of the transcriptional activation signalling GB_SynP promoters was also demonstrated, since co-expression of dCasEV2.1 with an unrelated gRNA led to negligible basal expression in all assays. These results position the GB_SynP collection as a promising tool for the regulation of complex multigene circuits with different gRNAs present in the cis-regulatory boxes of each promoter, thus creating logic gates that could be useful to further explore different metabolic fluxes within biosynthetic pathways. Moreover, other studies reported the successful expression of gRNAs under pol-II promoters, which in turn could be regulated by different inducers ^21,27^, thus allowing customizable control of each gRNA with different stimuli to further direct the multigene circuits in different ways.

While we reported here the assembly and behaviour of 35 synthetic promoters, the GB_SynP collection includes to date 32 promoter parts, compiled in Table S2, that can be used to assemble more than 500 different promoters without the need for creating any new sequence, standing out as one of the CRISPRa-based synthetic expression tools currently available for plants with the highest diversity and combinatorial strength. Moreover, plant synthetic promoters created so far mostly rely on the well-characterised CaMV *35s* minimal promoter ^27,30^, which might lead to higher basal expression in comparison with other minimal promoters like mDFR ^21^. To overcome this limitation, GB_SynP includes newly designed minimal promoter parts, for which negligible basal expression was shown in all cases, as well as a range of activation levels. The total length of the synthetic promoter should also be considered, as short sequences could easily be interfered with by other nearby promoters once they are introduced into the plant genome, especially considering the preference for T-DNA to be inserted into transcriptionally active regions ^31,32^. In this regard, different A1 random parts were also included in the GB_SynP collection to allow easy modulation of the length of the resulting promoter, while adding an extra source of sequence variation.

Although dCasEV2.1 remains as the most optimal system to regulate GB_SynP synthetic promoters, here we demonstrated that their activation can also be triggered by combining dCas9 and MS2 proteins with other activation domains, or by using other CRISPRa strategies, such as those based on dCas9:SunTag fusion. Among the combinations tested, the VPR activation domain showed the highest activation levels for both systems, which correlates with what was previously observed by Chavez et al. ^9^ where VPR reached the highest fluorescence values out of the 22 different activation domains tested, including the commonly used VP64. VPR is in fact a combination of the activation domains VP64, P65 and Rta, of which VP64 is in turn comprised of four tetrameric repetitions of the herpes simplex virus VP16 protein.

Depending on the intended application of GB_SynP promoters, concerns may arise from the use of viral proteins for the regulation of GB_SynP promoters. In this regard, as an alternative, we propose the combination of dCas9 and MS2 proteins with the EDLL and ERF2 activation domains, respectively, which triggered a considerable activation that led to a signal comparable to a NOS promoter level. Nevertheless, better-performing CRISPRa tools are continuously being developed ^12^ which could also be used in combination with different activation domains to increase the expression levels of GB_SynP promoters developed here.

The combinatorial power and the wide range of expression levels provided by the GB_SynP collection were further exploited in the optimisation of a multigene bioluminescence pathway. The new synthetic promoters were shown to regulate the expression three genes in the pathway in a predictable and reliable way, with the lowest pathway output levels (luminescence) obtained when all three genes were under the regulation of the weakest promoter, and the highest expression reached when the three genes were driven by the strongest promoters. In this case, we showed that the GB_SynP system was also useful to further characterise the regulatory requirements of the synthetic pathway. We found that, unlike the rest of the genes, low HispS expression limits the flux in the pathway, rendering the regulation of the remaining steps useless. Such behaviour is in line with previous observations described by Mitiouchkina et al. ^23^, where they reported that the addition of caffeic acid to *N. benthamiana* leaves expressing the auto-luminescence pathway resulted in the development of lower and slower luminescence than the addition of hispidin of luciferin.

Lucks et al. ^33^ defined five fundamental characteristics for efficient and predictable genetic engineering, which are independence, orthogonality, reliability, tunability and composability. Our GB_SynP system described here is a modular and composable system that has shown to be highly gRNA-specific and whose orthogonality is ensured by the negligible basal expression of the synthetic promoters generated when used in combination with the genome-wide specific dCasEV2.1 system ^18^. We further demonstrated that the GB_SynP system works in a reliable way for expressing the bioluminescence pathway and includes a range of expression levels that can be further modulated by the use of inducible pol-II driven gRNAs. All in all, the GB_SynP system constitutes a promising tool for the easy design and optimization of multigenic circuits in the field of plant genetic engineering.

## MATERIALS AND METHODS

### Construction and assembly of DNA parts

All plasmids used in this work were assembled using GoldenBraid (GB) cloning ^34^. The DNA sequences of the constructs generated in this work are available at https://gbcloning.upv.es/search/features by entering the IDs provided in Table S3. Random DNA sequences were generated at https://www.bioinformatics.org/sms2/random_dna.html ^35^ and each promoter part designed was ordered as gBlocks (IDT) and assembled following GoldenBraid (GB) domestication strategy. Briefly, DNA parts were first cloned into the pUPD2 entry vector and verified by digestion and sequencing. Transcriptional units were then generated via restriction-ligation reactions with the different DNA parts contained in pUPD2 vectors, and combined with binary assemblies into multigenic constructs via restriction-ligation with T4 ligase and BsaI or BsmBI. All constructs were cloned into *Escherichia coli* TOP 10 strain using Mix&Go kit (Zymo Research) as indicated by the manufacturer. All assemblies were confirmed by digestion.

### Plant inoculation and transient expression assays

Transient expression assays were performed by agroinfiltration of 4-5-week-old *N. benthamiana* plants grown at 24°C/20°C (light/darkness) with a 16h:8h photoperiod. Expression vectors were transferred to *Agrobacterium tumefaciens* GV3101 by electroporation. Cultures were grown overnight in liquid LB medium supplemented with rifampicin and the corresponding antibiotic for plasmid selection. Cells were then pelleted and resuspended in agroinfiltration buffer (10 mM MES at pH 5.6, 10 mM MgCl_2_ and 200 μM acetosyringone), incubated for 2 hours in the dark, and adjusted to an OD_600_ of 0.1. For co-infiltration, cultures were mixed at equal volumes. The silencing suppressor P19 was included in all tested constructs. Agroinfiltration was carried out using 1 mL needleless syringe through the abaxial surface of the three youngest fully expanded leaves.

### *In vitro* Luciferase/Renilla assay

Agroinfiltrated samples were collected 5 days post-infiltration using a Ø 8 mm corkborer and snap frozen in liquid nitrogen. Expression of Firefly luciferase (FLuc) and Renilla luciferase (RLuc) were determined with the Dual-Glo® Luciferase Assay System (Promega) following manufacturer’s instruction with some modifications. Frozen leaf samples were first homogenized and extracted with 180µl Passive Lysis Buffer, followed by a centrifugation (14,000×g) at 4 °C for 10 minutes. 10 µl of working plant extract (supernatant) was then transferred to a 96 wellplate, where 40 µl LARII buffer was added to measure Fluc signal in a GloMax 96 Microplate Luminometer (Promega) with a 2-s delay and a 10-s measurement. RLuc signal was measured afterwards by adding 40 µl Stop&Glow reagent and measuring in the same way.

FLuc/RLuc ratios were determined as the mean value of three independent agroinfiltrated leaves of the same plant and were normalized to the FLuc/RLuc ratio obtained from a sample agroinfiltrated with a reference construct (GB1398) where Luciferase is driven by *NOS* promoter (p*NOS*) and Renilla is under CaMV *35s* promoter (p*35s*). Reference FLuc/RLuc ratios are arbitrarily set as 1.0 relative promoter units (RPUs).

### *In vivo* Luciferase/eGFP assay

Agroinfiltrated leaf discs were collected 24h post-infiltration using a Ø 6 mm corkborer, and placed directly in 96 wellplates containing 200 µL/well of solid MS medium (4.9 g/L MS+vitamins, 8 g/L agar pH=5.7). Plates were measured once per day for 8 days in a GloMax 96 Microplate Luminometer (Promega). For luminescence, a 2-s delay and 10-s measurement parameters were used as previously described for in vitro Luciferase/Renilla assays. For eGFP measurement, an optical kit was used with an excitation peak at 490nm and emission at 510-570 nm.

## ABBREVIATIONS

TF: transcription factor
GB: GoldenBraid
CRISPRa: CRISPR activation
dCas9: nuclease-deactivated Cas9 protein
NbDFR: NADPH-dependent dihydroflavonol-4-reductase gene from *Nicotiana benthamiana*
SlDFR: NADPH-dependent dihydroflavonol-4-reductase gene from *Solanum lycopersicum*
TSS: transcriptional start site
gRNA: guide RNA
dCasEV2.1: dCas9:EDLL + MS2:VPR + gRNA
pNOS: Nopaline synthase promoter
CaMV 35s: Cauliflower mosaic virus 35s gene
FLuc: Firefly luciferase
RLuc: Renilla luciferase
RPU: relative promoter unit
ScFv: single-chain variable fragment
eGFP: enhanced GFP protein

## AUTHOR INFORMATION

### Author contributions

E.M.-G., S.S., C.C. and D.O. designed the experiments. E.M.-G. and S.S. conducted the experiments. E.M.-G. and D.O. drafted the manuscript. All the authors revised the manuscript.

### SUPPORTING INFORMATION

Supplementary material: TableS1: Sequences of the (A3) minimal promoters designed in this study. TableS2: Summary of DNA parts included to date in the GB_SynP collection. TableS3: GoldenBraid DNA parts and assemblies created and/or used in this study. Figure S1: Alignment of A2 parts sequences included in the GB_SynP collection. Figure S2. Expression range of the GB_SynP promoters used for regulation of LUZ pathway. Figure S3. Basal expression of the Time-course experiment including the 27 constructs expressing the LUZ pathway transiently in *Nicotiana benthamiana* leaves under the regulation of 1x, 2x or 3x GB_SynP promoters.

## ACKNOWLEDGMENTS

This work was funded by Era-CoBiotech SUSPHIRE (PCI2018-092893) and PID2019-108203RB-100 Plan Nacional I+D, Spanish Ministry of Economy and Competitiveness. E.M.-G. and S.S. acknowledges support by a FPU (FPU18/02019) and a FPI (BIO2016-78601-R) H2020 Research Program: 760331 Newcotiana fellowships, respectively, from the Spanish Ministry of Science, Innovation and Universities. C.C. acknowledges support by a FPI-UPV (PAID-01-20) fellowship from Universitat Politècnica de València. The authors would like to thank Dr. Lynne Yenush for her assistance with the manuscript preparation.

## REFERENCES

1. Weber, E., Engler, C., Gruetzner, R., Werner, S., and Marillonnet, S. (2011) A modular cloning system for standardized assembly of multigene constructs. PLoS ONE 6.

2. Engler, C., Youles, M., Gruetzner, R., Ehnert, T. M., Werner, S., Jones, J. D. G., Patron, N. J., and Marillonnet, S. (2014) A Golden Gate modular cloning toolbox for plants. ACS Synthetic Biology 3, 839–843.

3. Sarrion-Perdigones, A., Falconi, E. E., Zandalinas, S. I., Juárez, P., Fernández-del-Carmen, A., Granell, A., and Orzaez, D. (2011) GoldenBraid: An iterative cloning system for standardized assembly of reusable genetic modules. PLoS ONE 6.

4. Andreou, A. I., and Nakayama, N. (2018) Mobius assembly: A versatile golden-gate framework towards universal DNA assembly. PLoS ONE 13.

5. Pollak, B., Cerda, A., Delmans, M., Álamos, S., Moyano, T., West, A., Gutiérrez, R. A., Patron, N. J., Federici, F., and Haseloff, J. (2019) Loop assembly: a simple and open system for recursive fabrication of DNA circuits. New Phytologist 222, 628–640.

6. Patron, N. J., Orzaez, D., Marillonnet, S., Warzecha, H., Matthewman, C., Youles, M., Raitskin, O., Leveau, A., Farré, G., Rogers, C., Smith, A., Hibberd, J., Webb, A. A. R., Locke, J., Schornack, S., Ajioka, J., Baulcombe, D. C., Zipfel, C., Kamoun, S., Jones, J. D. G., Kuhn, H., Robatzek, S., van Esse, H. P., Sanders, D., Oldroyd, G., Martin, C., Field, R., O’Connor, S., Fox, S., Wulff, B., Miller, B., Breakspear, A., Radhakrishnan, G., Delaux, P. M., Loqué, D., Granell, A., Tissier, A., Shih, P., Brutnell, T. P., Quick, W. P., Rischer, H., Fraser, P. D., Aharoni, A., Raines, C., South, P. F., Ané, J. M., Hamberger, B. R., Langdale, J., Stougaard, J., Bouwmeester, H., Udvardi, M., Murray, J. A. H., Ntoukakis, V., Schäfer, P., Denby, K., Edwards, K. J., Osbourn, A., and Haseloff, J. (2015) Standards for plant synthetic biology: a common syntax for exchange of DNA parts. New Phytologist 208, 13–19.

7. Cai, Y. M., Carrasco Lopez, J. A., and Patron, N. J. (2020) Phytobricks: Manual and Automated Assembly of Constructs for Engineering Plants. Methods Mol Biol 2205, 179–199.

8. Vazquez-Vilar, M., Quijano-Rubio, A., Fernandez-Del-Carmen, A., Sarrion-Perdigones, A., Ochoa-Fernandez, R., Ziarsolo, P., Jos’, J., Blanca, J., Granell, A., and Orzaez, D. (2017) GB3.0: a platform for plant bio-design that connects functional DNA elements with associated biological data. Nucleic Acids Research 45, 2196–2209.

9. Chavez, A., Scheiman, J., Vora, S., Pruitt, B. W., Tuttle, M., Iyer, P. R., Lin, S., Kiani, S., Guzman, C. D., Wiegand, D. J., Ter-Ovanesyan, D., Braff, J. L., Davidsohn, N., Housden, B. E., Perrimon, N., Weiss, R., Aach, J., Collins, J. J., and Church, G. M. (2015) Highly efficient cas9-mediated transcriptional programming. Nature Methods 326.

10. Li, Z., Zhang, D., Xiong, X., Yan, B., Xie, W., Sheen, J., and Li, J.-F. (2017) A potent Cas9-derived gene activator for plant and mammalian cells. Nat Plants 3, 930–936.

11. Lowder, L. G., Zhou, J., Zhang, Y., Malzahn, A., Zhong, Z., Hsieh, T. F., Voytas, D. F., Zhang, Y., and Qi, Y. (2018) Robust Transcriptional Activation in Plants Using Multiplexed CR.ISPR-Act2.0 and mTALE-Act Systems. Molecular Plant 11, 245–256.

12. Pan, C., Wu, X., Markel, K., Malzahn, A. A., Kundagrami, N., Sretenovic, S., Zhang, Y., Cheng, Y., Shih, P. M., and Qi, Y. (2021) CRISPR–Act3.0 for highly efficient multiplexed gene activation in plants. Nature Plants 2021 7:7 7, 942–953.

13. Vazquez-Vilar, M., Bernabé-Orts, J. M., Fernandez-del-Carmen, A., Ziarsolo, P., Blanca, J., Granell, A., and Orzaez, D. (2016) A modular toolbox for gRNA-Cas9 genome engineering in plants based on the GoldenBraid standard. Plant Methods 12, 1–12.

14. Papikian, A., Liu, W., Gallego-Bartolomé, J., and Jacobsen, S. E. (2019) Site-specific manipulation of Arabidopsis loci using CRISPR-Cas9 SunTag systems. Nature Communications.

15. Xiong, X., Liang, J., Li, Z., Gong, B. Q., and Li, J. F. (2021) Multiplex and optimization of dCas9-TV-mediated gene activation in plants. Journal of Integrative Plant Biology 63, 634–645.

16. Konermann, S., Brigham, M. D., Trevino, A. E., Joung, J., Abudayyeh, O. O., Barcena, C., Hsu, P. D., Habib, N., Gootenberg, J. S., Nishimasu, H., Nureki, O., and Zhang, F. (2015) Genome-scale transcriptional activation by an engineered CRISPR-Cas9 complex. Nature 517, 583–588.

17. Mali, P., Esvelt, K. M., and Church, G. M. (2013) Cas9 as a versatile tool for engineering biology. Nat Methods 10, 957–963.

18. Selma, S., Bernabé-Orts, J. M., Vazquez-Vilar, M., Diego-Martin, B., Ajenjo, M., Garcia-Carpintero, V., Granell, A., and Orzaez, D. (2019) Strong gene activation in plants with genome-wide specificity using a new orthogonal CRISPR/Cas9-based programmable transcriptional activator. Plant Biotechnology Journal. Blackwell Publishing Ltd.

19. Polstein, L. R., Perez-Pinera, P., Kocak, D. D., Vockley, C. M., Bledsoe, P., Song, L., Safi, A., Crawford, G. E., Reddy, T. E., and Gersbach, C. A. (2015) Genome-wide specificity of DNA binding, gene regulation, and chromatin remodeling by TALE-and CRISPR/Cas9-based transcriptional activators. Genome Research 25, 1158–1169.

20. Khakhar, A., Starker, C. G., Chamness, J. C., Lee, N., Stokke, S., Wang, C., Swanson, R., Rizvi, F., Imaizumi, T., and Voytas, D. F. (2020) Building customizable auto-luminescent luciferase-based reporters in plants. Elife 9.

21. Garcia-Perez, E., Diego-Martin, B., Quijano-Rubio, A., Moreno-Giménez, E., Selma, S., Orzaez, D., and Vazquez-Vilar, M. (2022) A copper switch for inducing CRISPR/Cas9-based transcriptional activation tightly regulates gene expression in Nicotiana benthamiana. BMC Biotechnology 2022 22:1 22, 1–13.

22. Kotlobay, A. A., Sarkisyan, K. S., Mokrushina, Y. A., Marcet-Houben, M., Serebrovskaya, E. O., Markina, N. M., Somermeyer, L. G., Gorokhovatsky, A. Y., Vvedensky, A., Purtov, K. v., Petushkov, V. N., Rodionova, N. S., Chepurnyh, T. v., Fakhranurova, L. I., Guglya, E. B., Ziganshin, R., Tsarkova, A. S., Kaskova, Z. M., Shender, V., Abakumov, M., Abakumova, T. O., Povolotskaya, I. S., Eroshkin, F. M., Zaraisky, A. G., Mishin, A. S., Dolgov, S. v., Mitiouchkina, T. Y., Kopantzev, E. P., Waldenmaier, H. E., Oliveira, A. G., Oba, Y., Barsova, E., Bogdanova, E. A., Gabaldón, T., Stevani, C. v., Lukyanov, S., Smirnov, I. v., Gitelson, J. I., Kondrashov, F. A., and Yampolsky, I. v. (2018) Genetically encodable bioluminescent system from fungi. Proc Natl Acad Sci U S A 115, 12728–12732.

23. Mitiouchkina, T., Mishin, A. S., Somermeyer, L. G., Markina, N. M., Chepurnyh, T. v., Guglya, E. B., Karataeva, T. A., Palkina, K. A., Shakhova, E. S., Fakhranurova, L. I., Chekova, S. v., Tsarkova, A. S., Golubev, Y. v., Negrebetsky, V. v., Dolgushin, S. A., Shalaev, P. v., Shlykov, D., Melnik, O. A., Shipunova, V. O., Deyev, S. M., Bubyrev, A. I., Pushin, A. S., Choob, V. v., Dolgov, S. v., Kondrashov, F. A., Yampolsky, I. v., and Sarkisyan, K. S. (2020) Plants with genetically encoded autoluminescence. Nature Biotechnology 38, 944–946.

24. Dong, C., Fontana, J., Patel, A., Carothers, J. M., and Zalatan, J. G. (2018) Synthetic CRISPR-Cas gene activators for transcriptional reprogramming in bacteria. Nature Communications 2018 9:1 9, 1–11.

25. Farzadfard, F., Perli, S. D., and Lu, T. K. (2013) Tunable and multifunctional eukaryotic transcription factors based on CRISPR/Cas. ACS Synthetic Biology 2, 604–613.

26. Nissim, L., Perli, S. D., Fridkin, A., Perez-Pinera, P., and Lu, T. K. (2014) Multiplexed and Programmable Regulation of Gene Networks with an Integrated RNA and CRISPR/Cas Toolkit in Human Cells. Molecular Cell 54, 698–710.

27. Kar, S., Bordiya, Y., Rodriguez, N., Kim, J., Gardner, E. C., Gollihar, J. D., Sung, S., and Ellington, A. D. (2022) Orthogonal control of gene expression in plants using synthetic promoters and CRISPR-based transcription factors. Plant Methods 2022 18:1 18, 1–15.

28. Morbitzer, R., Römer, P., Boch, J., and Lahaye, T. (2010) Regulation of selected genome loci using de novo-engineered transcription activator-like effector (TALE)-type transcription factors. Proc Natl Acad Sci U S A 107, 21617–21622.

29. Urnov, F. D., Rebar, E. J., Holmes, M. C., Zhang, H. S., and Gregory, P. D. (2010) Genome editing with engineered zinc finger nucleases. Nature Reviews Genetics 2010 11:9 11, 636–646.

30. Dey, N., Sarkar, S., Acharya, S., and Maiti, I. B. (2015) Synthetic promoters in planta. Planta 2015 242:5 242, 1077–1094.

31. Ingelbrecht, I., Breyne, P., Vancompernolle, K., Jacobs, A., van Montagu, M., and Depicker, A. (1991) Transcriptional interference in transgenic plants (Agrobacterium vector system; callus; position effect; poly(A) signal; recombinant DNA; terminator). Gene 109, 239–242.

32. Schneeberger, R. G., Zhang, K., Tatarinova, T., Troukhan, M., Kwok, S. F., Drais, J., Klinger, K., Orejudos, F., Macy, K., Bhakta, A., Burns, J., Subramanian, G., Donson, J., Flavell, R., and Feldmann, K. A. (2005) Agrobacterium T-DNA integration in Arabidopsis is correlated with DNA sequence compositions that occur frequently in gene promoter regions. Functional and Integrative Genomics 5, 240–253.

33. Lucks, J. B., Qi, L., Whitaker, W. R., and Arkin, A. P. (2008) Toward scalable parts families for predictable design of biological circuits. Current Opinion in Microbiology 11, 567–573.

34. Sarrion-Perdigones, A., Vazquez-Vilar, M., Palací, J., Castelijns, B., Forment, J., Ziarsolo, P., Blanca, J., Granell, A., and Orzaez, D. (2013) GoldenBraid 2.0: a comprehensive DNA assembly framework for plant synthetic biology. Plant Physiol 162, 1618–1631.

35. Stothard, P. (2000) The sequence manipulation suite: JavaScript programs for analyzing and formatting protein and DNA sequences. Biotechniques 28.

