## Supplementary material for "GB_SynP: a modular dCas9-regulated synthetic promoter collection for fine-tuned recombinant gene expression in plants"

**TableS1. Sequences of the A3 minimal promoters designed in this study for the GB_SynP collection.** Highlighted in bold are the predicted TATA boxes of each promoter sequence. Underlined sequences corresponds to predicted 5' UTR regions.

| GB CODE | PART NAME | ORGANISM | SEQUENCE (TATA BOX IN BOLD, 5' UTR UNDERLINED) |
|---|---|---|---|
| **GB3408** | mCHS | *Solanum lycopersicum* | atctctagctaccattctttttcttttggacttca**tatataaa**tatatagttcatacaaccccatgcaacaaaaatacaccaagacaattatcactctttcattcacgtagtcctaaacacaaaaaaacctagcatatccaccatttttccggcgaaa |
| **GB3409** | mANS | *Solanum lycopersicum* | tatcttgtatgagataaatttcaagtgc**tataaa**tatacctaacgaaaagttatagttagcagtactgaaaaaacgagataaac |
| **GB3410** | mTA29 | *Nicotiana tabacum* | ttttgcaagttgttgtgtatgtcttgtgctc**tatata**tgcccttgtggtgcaagtgtaacagtacaacatcatcactcaaatcaaagtttttacttaaagaaattagctaaa |
| **GB3412** | mPCPS2 | *Nicotiana tabacum* | tttaatatttgcccttcatcaagaggcaata**tataaa**taagcctgcgccattgcaacaactcaaaccatttccaatattgcctcacaagtcagtagtgcctttcctctctcaaacgttcattgtctttatctcccttccccaattctcattggaagaataaaacaaaaattaaattagaa |
| **GB3411** | m2S3 | *Arabidopsis thaliana* | gcatgcatgcattcttacacgtgattgccatgcaaatctcctttctcacc**tataaa**tacaaaccaacccttcactacactcttcactcaaaccaaaacaagaaaacatacacaaatagcaaaac |
| **GB3413** | mPAF | *Penicillium chrysogenum* | tgtcacctttcacacagagctcatgatctggtt**tataa**aggcggcttcatgaccctcaattccatatagtatcactcccatcacagcatttcgatatcttcaaccactttaaccttctccagaggatcatcatctcaagcccttcata |
| **GB3414** | mGPDA | *Aspergillus nidulans* | tttgttgcca**tatttt**cctgctctccccaccagctgctcttttcttttctctttcttttcccatcttcagtatattcatcttcccatccaagaacctttaa |

**TableS2. Summary of all DNA parts included to date in the GB_SynP collection.**

| GB_SynP A1 parts (distal promoters) | GB_SynP A2 parts (proximal promoters) | | | | | GB_SynP A3 parts (minimal promoters) |
|---|---|---|---|---|---|---|
| | 1x target for gRNA1 | 2x target for gRNA1 | 3x target for gRNA1 | 1 target for gRNA2 | 1 target for gRNA3 | |
| *R1* | G1aG2b.1 | G1ab.1 | G1abc.1 | G1aG2b.1 | G3aG2b.1 | mDFR |
| *R2* | G1aG2b.2 | G1ab.2 | G1abc.2 | G1aG2b.2 | | mANS |
| *R3* | G1aG2b.3 | G1ab.3 | G1abc.3 | G1aG2b.3 | | mCHS |
| | G1aG2b.4 | G1ab.4 | G1abc.4 | G1aG2b.4 | | mPCPS2 |
| | G1aG2b.5 | G1ab.5 | G1abc.5 | G1aG2b.5 | | mTA29 |
| | G1aG2b.6 | | | G1aG2b.6 | | m2S3 |
| | G1aG2b.7 | | | G1aG2b.7 | | mPAF |
| | G1dG2b.1 | | | G1dG2b.1 | | mGPDA |
| | G1b.1 | | | | | |
| | G1e.1 | | | | | |

**TableS3. DNA parts and assemblies created and/or used in this study.**

| GB Code | Part Name | Description |
|---|---|---|
| GB0164 | pEGB 35s:Luciferase:TNos-SF-35s:Renilla:TNos-35s:P19:TNos | Module for the expression of the Firefly luciferase, the Renilla luciferasa and the P19 silencing suppressor genes driven by the 35s promoter and the nos terminator. |
| GB1160 | pEGB SlDFR:Luc:TNos-SF-35S:Renilla:TNos-35S:P19:TNos | Module for the expression of the Firefly luciferase gene driven by the S.lycopersicum DFR promoter and terminator, the Renilla luciferase gene and the silencing suppressor P19 driven by the 35s promoter and the Nos terminator. |
| GB1190 | pEGB 35s:dCas9-EDLL:tNOS | TU for the expression of the dCas9 with the activation domain EDLL as a CT fusion. |
| GB1398 | pEGB3alpha2 Pnos:luc:Tnos-SF-35S:Ren:Tnos-35s:P19:Tnos-SF | Module for the expression of the Firefly Luciferase under the regulation of the Nos promoter, and the Renilla Luciferase and the P19 silencing suppressor under the regulation of the 35S promoter. |
| GB1603 | 35s:dCas9-SunTag:Tnos | TU for the expression of the dCas9 fused to the SunTag epitope. |
| GB1738 | Alpha2: 35s_MS2:EDLL_Tnos | TU for the constitutive expression of the MS2 coat protein fused on Ct to the activation domain EDLL. |
| GB1824 | Alpha2_35s-dCas9:ERF2-Tnos | TU for the constitutive expression of dCas9 fused to the transcriptional activation domain ERF2. |
| GB1833 | Alpha2_35s-MS2:ERF2-Tnos | TU for the constitutive expression of the MS2 coat protein fused on Ct to the transactivation domain of ERF2. |
| GB1867 | Alpha2_35s-ScFv:VPR-Tnos | TU for constitutive expression of ScFv antibody fused to VPR activation. |
| GB1869 | Alpha2_35s-ScFv:ERF2-Tnos | TU for constitutive expression of ScFv antibody fused to ERF2 activation domain. |
| GB2085 | Omega1_35s-Ms2:VPR-Tnos-35s-dCas9:EDLL-Tnos | Module for the expression of Ms2 protein fused to VPR and dCas9 fused to EDLL. |
| GB2034 | Alpha1: 2gRNA SlDFR | Multiplex construct to target SlDFR in -150 position. |
| GB2035 | Alpha1: 3gRNA SlDFR | Multiplex construct to target SlDFR in +31 position. |
| GB2036 | Alpha1: 4gRNA SlDFR | Multiplex construct to target SlDFR in -1320 position. |
| GB2037 | Alpha1:5gRNA SlDFR | Multiplex construct to target SlDFR in -1052 position. |
| GB2038 | Alpha1:6gRNA SlDFR | Multiplex construct to target SlDFR in -979 position. |
| GB2039 | Alpha1:7gRNA SlDFR | Multiplex construct to target SlDFR in -683 position. |
| GB2566 | pUPD2_miniDFR | Minimal promoter of SlDFR gene containing 62bp upstream the transcription start site and the 5'UTR region. |
| GB2815 | pUPD2_GB_SynP (A1) Random Sequence R1 | Random sequence R1 of 1240 bp for A1 distal promoter position. |
| GB2878 | pUPD2_GB_SynP (A2) G1aG2b.1 | A2 Proximal promoter sequence consisting of the target sequences for the gRNA-1 DFR (gRNA1) and gRNA-5 DFR (gRNA2) flanked by random sequences. |
| GB2879 | pUPD2_GB_SynP (A2) G1aG2b.5 | A2 Proximal promoter sequence consisting of the target sequences for the gRNA-1 DFR (gRNA1) and gRNA-5 DFR (gRNA2) flanked by random sequences. |
| GB2880 | pUPD2_GB_SynP (A2) G1aG2b.6 | A2 Proximal promoter sequence consisting of the target sequences for the gRNA-1 DFR (gRNA1) and gRNA-5 DFR (gRNA2) flanked by random sequences. |
| GB2881 | pUPD2_GB_SynP (A2) G1aG2b.2 | A2 Proximal promoter sequence consisting of the target sequences for the gRNA-1 DFR (gRNA1) and gRNA-5 DFR (gRNA2) flanked by random sequences. |
| GB2882 | pUPD2_GB_SynP (A2) G1aG2b.3 | A2 Proximal promoter sequence consisting of the target sequences for the gRNA-1 DFR (gRNA1) and gRNA-5 DFR (gRNA2) flanked by random sequences. |
| GB2883 | pUPD2_GB_SynP (A2) G1aG2b.4 | A2 Proximal promoter sequence consisting of the target sequences for the gRNA-1 DFR (gRNA1) and gRNA-5 DFR (gRNA2) flanked by random sequences. |
| GB2884 | pUPD2_GB_SynP (A2) G1ab.2 | A2 Proximal promoter sequence consisting of two times the target sequence for the gRNA-1 DFR (gRNA1) flanked by random sequences. |
| GB2885 | pUPD2_GB_SynP (A2) G1ab.1 | A2 Proximal promoter sequence consisting of two times the target sequence for the gRNA-1 DFR (gRNA1) flanked by random sequences. |

| | | |
|---|---|---|
| GB2886 | pUPD2_GB_SynP (A2) G1ab.3 | A2 Proximal promoter sequence consisting of two times the target sequence for the gRNA-1 DFR (gRNA1) flanked by random sequences. |
| GB2887 | pUPD2_GB_SynP (A2) G1ab.6 | A2 Proximal promoter sequence consisting of two times the target sequence for the gRNA-1 DFR (gRNA1) flanked by random sequences. |
| GB2888 | pUPD2_GB_SynP (A2) G1ab.4 | A2 Proximal promoter sequence consisting of two times the target sequence for the gRNA-1 DFR (gRNA1) flanked by random sequences. |
| GB2889 | pUPD2_GB_SynP (A2) G1ab.5 | A2 Proximal promoter sequence consisting of two times the target sequence for the gRNA-1 DFR (gRNA1) flanked by random sequences. |
| GB2890 | pUPD2_GB_SynP (A2) G1dG2b.1 | A2 Proximal promoter sequence consisting of the target sequence for the gRNA-1 DFR (gRNA1, at "d site" positioned at -120 from TSS) and gRNA-5 DFR (gRNA2) flanked by random sequences. |
| GB2891 | pUPD2_GB_SynP (A2) G1abc.1 | A2 Proximal promoter sequence consisting of three times the target sequence for the gRNA-1 DFR (gRNA1) flanked by random sequences. |
| GB2894 | Alpha1_R1:G1aG2b.1:mDFR:luc:t35s | TU for the expression of firefly luciferase driven by a GB_SynP promoter with A1 R1 and A3 mSlDFR parts, and an A2 part containing a target for the gRNA-1DFR (gRNA1) and gRNA-5 DFR (gRNA2). |
| GB2895 | Alpha1_R1:G1aG2b.5:mDFR:luc:t35s | TU for the expression of firefly luciferase driven by a GB_SynP promoter with A1 R1 and A3 mSlDFR parts, and an A2 part containing a target for the gRNA-1DFR (gRNA1) and gRNA-5 DFR (gRNA2). |
| GB2896 | Alpha1_R1:G1aG2b.6:mDFR:luc:t35s | TU for the expression of firefly luciferase driven by a GB_SynP promoter with A1 R1 and A3 mSlDFR parts, and an A2 part containing a target for the gRNA-1DFR (gRNA1) and gRNA-5 DFR (gRNA2). |
| GB2897 | Alpha1_R1:G1aG2b.2:mDFR:luc:t35s | TU for the expression of firefly luciferase driven by a GB_SynP promoter with A1 R1 and A3 mSlDFR parts, and an A2 part containing a target for the gRNA-1DFR (gRNA1) and gRNA-5 DFR (gRNA2). |
| GB2898 | Alpha1_R1:G1aG2b.3:mDFR:luc:t35s | TU for the expression of firefly luciferase driven by a GB_SynP promoter with A1 R1 and A3 mSlDFR parts, and an A2 part containing a target for the gRNA-1DFR (gRNA1) and gRNA-5 DFR (gRNA2). |
| GB2899 | Alpha1_R1:G1aG2b.4:mDFR:luc:t35s | TU for the expression of firefly luciferase driven by a GB_SynP promoter with A1 R1 and A3 mSlDFR parts, and an A2 part containing a target for the gRNA-1DFR (gRNA1) and gRNA-5 DFR (gRNA2). |
| GB2900 | Alpha1_R1:G1ab.2:mDFR:luc:t35s | TU for the expression of firefly luciferase driven by a GB_SynP promoter with A1 R1 and A3 mSlDFR parts, and an A2 part containing the target for the gRNA-1DFR (gRNA1) twice. |
| GB2901 | Alpha1_R1:G1ab.1:mDFR:luc:t35s | TU for the expression of firefly luciferase driven by a GB_SynP promoter with A1 R1 and A3 mSlDFR parts, and an A2 part containing the target for the gRNA-1DFR (gRNA1) twice. |
| GB2902 | Alpha1_R1:G1ab.3:mDFR:luc:t35s | TU for the expression of firefly luciferase driven by a GB_SynP promoter with A1 R1 and A3 mSlDFR parts, and an A2 part containing the target for the gRNA-1DFR (gRNA1) twice. |
| GB2904 | Alpha1_R1:G1ab.4:mDFR:luc:t35s | TU for the expression of firefly luciferase driven by a GB_SynP promoter with A1 R1 and A3 mSlDFR parts, and an A2 part containing the target for the gRNA-1DFR (gRNA1) twice. |
| GB2905 | Alpha1_R1:G1ab.5:mDFR:luc:t35s | TU for the expression of firefly luciferase driven by a GB_SynP promoter with A1 R1 and A3 mSlDFR parts, and an A2 part containing the target for the gRNA-1DFR (gRNA1) twice. |
| GB2906 | Alpha1_R1:G1dG2b.1:mDFR:luc:t35s | TU for the expression of firefly luciferase driven by a GB_SynP promoter with A1 R1 and A3 mSlDFR parts, and an A2 part containing a target for the gRNA-1DFR (gRNA1, at "d site") and gRNA-5 DFR (gRNA2). |
| GB2907 | Alpha1_R1:G1abc.1:mDFR:luc:t35s | TU for the expression of firefly luciferase driven by a GB_SynP promoter with A1 R1 and A3 mSlDFR parts, and an A2 part containing three times the gRNA-1DFR (gRNA1) target sequence. |
| GB2909 | Omega1_R1:G1aG2b.1:mDFR:luc:t35s +P19(TU) +Renilla(TU) | Module for the expression of luciferase driven by a GB_SynP promoter with A1 R1, A3 mSlDFR, and an A2 part containing a target for gRNA1 and gRNA2, and for the constitutive expression of renilla and P19. |
| GB2910 | Omega1_R1:G1aG2b.5:mDFR:luc:t35s +P19(TU) +Renilla(TU) | Module for the expression of luciferase driven by a GB_SynP promoter with A1 R1, A3 mSlDFR, and an A2 part containing a target for gRNA1 and gRNA2, and for the constitutive expression of renilla and P19. |
| GB2912 | Omega1_R1:G1aG2b.2:mDFR:luc:t35s +P19(TU) +Renilla(TU) | Module for the expression of luciferase driven by a GB_SynP promoter with A1 R1, A3 mSlDFR, and an A2 part containing a target |

| | | |
|---|---|---|
| | | for gRNA1 and gRNA2, and for the constitutive expression of renilla and P19. |
| GB2913 | Omega1_R1:G1aG2b.3:mDFR:luc:t35s +P19(TU) +Renilla(TU) | Module for the expression of luciferase driven by a GB_SynP promoter with A1 R1, A3 mSlDFR, and an A2 part containing a target for gRNA1 and gRNA2, and for the constitutive expression of renilla and P19. |
| GB2914 | Omega1_R1:G1aG2b.4:mDFR:luc:t35s +P19(TU) +Renilla(TU) | Module for the expression of luciferase driven by a GB_SynP promoter with A1 R1, A3 mSlDFR, and an A2 part containing a target for gRNA1 and gRNA2, and for the constitutive expression of renilla and P19. |
| GB2915 | Omega1_R1:G1ab.2:mDFR:luc:t35s +P19(TU) +Renilla(TU) | Module for the expression of luciferase driven by a GB_SynP promoter with A1 R1, A3 mSlDFR, and an A2 part with the target for gRNA1 twice, and the constitutive expression of renilla and P19. |
| GB2916 | Omega1_R1:G1ab.1:mDFR:luc:t35s +P19(TU) +Renilla(TU) | Module for the expression of luciferase driven by a GB_SynP promoter with A1 R1, A3 mSlDFR, and an A2 part with the target for gRNA1 twice, and the constitutive expression of renilla and P19. |
| GB2917 | Omega1_R1:G1ab.3:mDFR:luc:t35s +P19(TU) +Renilla(TU) | Module for the expression of luciferase driven by a GB_SynP promoter with A1 R1, A3 mSlDFR, and an A2 part with the target for gRNA1 twice, and the constitutive expression of renilla and P19. |
| GB2919 | Omega1_R1:G1ab.4:mDFR:luc:t35s +P19(TU) +Renilla(TU) | Module for the expression of luciferase driven by a GB_SynP promoter with A1 R1, A3 mSlDFR, and an A2 part with the target for gRNA1 twice, and the constitutive expression of renilla and P19. |
| GB2920 | Omega1_R1:G1ab.5:mDFR:luc:t35s +P19(TU) +Renilla(TU) | Module for the expression of luciferase driven by a GB_SynP promoter with A1 R1, A3 mSlDFR, and an A2 part with the target for gRNA1 twice, and the constitutive expression of renilla and P19. |
| GB2921 | Omega1_R1:G1dG2b.1:mDFR:luc:t35s +P19(TU) +Renilla(TU) | Module for the expression of luciferase driven by a GB_SynP promoter with A1 R1, A3 mSlDFR, and an A2 part containing a target for gRNA1 (d site) and gRNA2, and for the constitutive expression of renilla and P19. |
| GB2922 | Omega1_R1:G1abc.1:mDFR:luc:t35s +P19(TU) +Renilla(TU) | Module for the expression of luciferase driven by a GB_SynP promoter with A1 R1, A3 mSlDFR, and an A2 part with three times the target for gRNA1, and the constitutive expression of renilla and P19. |
| GB3269 | pUPD2_GB_SynP (A1) Random Sequence R2 | Random sequence R2 of 1240 bp for A1 distal promoter position. |
| GB3270 | pUPD2_GB_SynP (A1) Random Sequence R3 | Random sequence R3 of 1240 bp for A1 distal promoter position. |
| GB3271 | pUPD2_GB_SynP (A2) G1aG2b.7 | A2 Proximal promoter sequence consisting of the target sequence for the gRNA-1 DFR (gRNA1) and gRNA-5 DFR (gRNA2) flanked by random sequences. |
| GB3272 | pUPD2_GB_SynP (A2) G1b.1 | A2 Proximal promoter sequence consisting of the target sequence for the gRNA-1 DFR (gRNA1) at "b site" (-210 from TSS). |
| GB3273 | pUPD2_GB_SynP (A2) G1e.1 | A2 Proximal promoter sequence consisting of the target sequence for the gRNA-1 DFR (gRNA1) at "e site" (-320 from TSS). |
| GB3274 | pUPD2_GB_SynP (A2) G3aG2b.1 | A2 Proximal promoter sequence consisting of the target sequences for the gRNA-4 NOS (gRNA3) and gRNA-5 DFR (gRNA2) flanked by random sequences. |
| GB3275 | pUPD2_GB_SynP (A2) G1abc.2 | A2 Proximal promoter sequence consisting of three times the target sequence for the gRNA-1 DFR (gRNA1) flanked by random sequences. |
| GB3276 | pUPD2_GB_SynP (A2) G1abc.3 | A2 Proximal promoter sequence consisting of three times the target sequence for the gRNA-1 DFR (gRNA1) flanked by random sequences. |
| GB3277 | pUPD2_GB_SynP (A2) G1abc.4 | A2 Proximal promoter sequence consisting of three times the target sequence for the gRNA-1 DFR (gRNA1) flanked by random sequences. |
| GB3278 | pUPD2_GB_SynP (A2) G1abc.5 | A2 Proximal promoter sequence consisting of three times the target sequence for the gRNA-1 DFR (gRNA1) flanked by random sequences. |
| GB3279 | Alpha1_R2:G1aG2b.1:mDFR:luc:t35s | TU for the expression of firefly luciferase driven by a GB_SynP promoter with A1 R2 and A3 mSlDFR parts, and an A2 part containing a target for the gRNA-1DFR (gRNA1) and gRNA-5 DFR (gRNA2). |
| GB3280 | Alpha1_R2:G1ab.1:mDFR:luc:t35s | TU for the expression of firefly luciferase driven by a GB_SynP promoter with A1 R2 and A3 mSlDFR parts, and an A2 part containing the target for the gRNA-1DFR (gRNA1) twice. |
| GB3281 | Alpha1_R2:G1abc:mDFR:luc:t35s | TU for the expression of firefly luciferase driven by a GB_SynP promoter with A1 R2 and A3 mSlDFR parts, and an A2 part containing three times the gRNA-1DFR (gRNA1) target sequence. |

| | | |
|---|---|---|
| GB3282 | Alpha1_R3:G1aG2b.1:mDFR:luc:t35s | TU for the expression of firefly luciferase driven by a GB_SynP promoter with A1 R3 and A3 mSlDFR parts, and an A2 part containing a target for the gRNA-1DFR (gRNA1) and gRNA-5 DFR (gRNA2). |
| GB3283 | Alpha1_R3:G1ab.1:mDFR:luc:t35s | TU for the expression of firefly luciferase driven by a GB_SynP promoter with A1 R3 and A3 mSlDFR parts, and an A2 part containing the target for the gRNA-1DFR (gRNA1) twice. |
| GB3284 | Alpha1_R3:G1abc:mDFR:luc:t35s | TU for the expression of firefly luciferase driven by a GB_SynP promoter with A1 R3 and A3 mSlDFR parts, and an A2 part containing three times the gRNA-1DFR (gRNA1) target sequence. |
| GB3285 | Omega1_R2:G1aG2b.1:mDFR:luc:t35s +P19(TU) +Renilla(TU) +P19(TU) +Renilla(TU) | Module for the expression of luciferase driven by a GB_SynP promoter with A1 R2, A3 mSlDFR, and an A2 part containing a target for gRNA1 and gRNA2, and for the constitutive expression of renilla and P19. |
| GB3286 | Omega1_R2:G1ab.1:mDFR:luc:t35s +P19(TU) +Renilla(TU) | Module for the expression of luciferase driven by a GB_SynP promoter with A1 R2, A3 mSlDFR, and an A2 part with the target for gRNA1 twice, and the constitutive expression of renilla and P19. |
| GB3287 | Omega1_R2:G1abc:mDFR:luc:t35s +P19(TU) +Renilla(TU) | Module for the expression of luciferase driven by a GB_SynP promoter with A1 R2, A3 mSlDFR, and an A2 part with three times the target for gRNA1, and the constitutive expression of renilla and P19. |
| GB3288 | Omega1_R3:G1aG2b.1:mDFR:luc:t35s +P19(TU) +Renilla(TU) | Module for the expression of luciferase driven by a GB_SynP promoter with A1 R3, A3 mSlDFR, and an A2 part containing a target for gRNA1 and gRNA2, and for the constitutive expression of renilla and P19. |
| GB3289 | Omega1_R3:G1ab.1:mDFR:luc:t35s +P19(TU) +Renilla(TU) | Module for the expression of luciferase driven by a GB_SynP promoter with A1 R3, A3 mSlDFR, and an A2 part with the target for gRNA1 twice, and the constitutive expression of renilla and P19. |
| GB3290 | Omega1_R3:G1abc:mDFR:luc:t35s +P19(TU) +Renilla(TU) | Module for the expression of luciferase driven by a GB_SynP promoter with A1 R3, A3 mSlDFR, and an A2 part with three times the target for gRNA1, and the constitutive expression of renilla and P19. |
| GB3317 | Alpha1_R1:G1aG2b.7:mDFR:luc:t35s | TU for the expression of firefly luciferase driven by a GB_SynP promoter with A1 R1 and A3 mSlDFR parts, and an A2 part containing a target for the gRNA-1DFR (gRNA1) and gRNA-5 DFR (gRNA2). |
| GB3318 | Alpha1_R1:G1b.1:mDFR:luc:t35s | TU for the expression of firefly luciferase driven by a GB_SynP promoter with A1 R1 and A3 mSlDFR parts, and an A2 part containing a target for the gRNA-1DFR (gRNA1, at "b site"). |
| GB3319 | Alpha1_R1:G1e.1:mDFR:luc:t35s | TU for the expression of firefly luciferase driven by a GB_SynP promoter with A1 R1 and A3 mSlDFR parts, and an A2 part containing a target for the gRNA-1DFR (gRNA1, at "e site"). |
| GB3320 | Alpha1_R1:G3aG2b.1:mDFR:luc:t35s | TU for the expression of firefly luciferase driven by a GB_SynP promoter with A1 R1 and A3 mSlDFR parts, and an A2 part containing a target for the gRNA-4 NOS (gRNA3) and gRNA-5 DFR (gRNA2). |
| GB3321 | Alpha1_R1:G1abc.2:mDFR:luc:t35s | TU for the expression of luciferase driven by a GB_SynP promoter with A1 R1 and A3 mSlDFR, and an A2 part with three times the gRNA1-DFR (gRNA1) target sequence. |
| GB3322 | Alpha1_R1:G1abc.3:mDFR:luc:t35s | TU for the expression of luciferase driven by a GB_SynP promoter with A1 R1 and A3 mSlDFR, and an A2 part with three times the gRNA1-DFR (gRNA1) target sequence. |
| GB3323 | Alpha1_R1:G1abc.4:mDFR:luc:t35s | TU for the expression of luciferase driven by a GB_SynP promoter with A1 R1 and A3 mSlDFR, and an A2 part with three times the gRNA1-DFR (gRNA1) target sequence. |
| GB3324 | Alpha1_R1:G1abc.5:mDFR:luc:t35s | TU for the expression of luciferase driven by a GB_SynP promoter with A1 R1 and A3 mSlDFR, and an A2 part with three times the gRNA1-DFR (gRNA1) target sequence. |
| GB3325 | Omega1_R1:G1aG2b.7:mDFR:luc:t35s +P19(TU) +Renilla(TU) | Module for the expression of luciferase driven by a GB_SynP promoter with A1 R1, A3 mSlDFR, and an A2 part containing a target for gRNA1 and gRNA2, and for the constitutive expression of renilla and P19. |
| GB3326 | Omega1_R1:G1b.1:mDFR:luc:t35s +P19(TU) +Renilla(TU) | Module for the expression of luciferase driven by a GB_SynP promoter with A1 R1, A3 mSlDFR, and an A2 part containing a target for gRNA1 (b site) and gRNA2, and for the constitutive expression of renilla and P19. |
| GB3327 | Omega1_R1:G1e.1:mDFR:luc:t35s +P19(TU) +Renilla(TU) | Module for the expression of luciferase driven by a GB_SynP promoter with A1 R1, A3 mSlDFR, and an A2 part containing a target for gRNA1 (e site) and gRNA2, and for the constitutive expression of renilla and P19. |

| | | |
|---|---|---|
| GB3328 | Omega1_R1:G3aG2b.1:mDFR:luc:t35s +P19(TU) +Renilla(TU) | Module for the expression of luciferase driven by a GB_SynP promoter with A1 R1, A3 mSlDFR, and an A2 part containing a target for gRNA3 and gRNA2, and for the constitutive expression of renilla and P19. |
| GB3329 | Omega1_R1:G1abc.2:mDFR:luc:t35s +P19(TU) +Renilla(TU) | Module for the expression of luciferase driven by a GB_SynP promoter with A1 R1, A3 mSlDFR, and an A2 part with three times the target for gRNA1, and the constitutive expression of renilla and P19. |
| GB3330 | Omega1_R1:G1abc.3:mDFR:luc:t35s +P19(TU) +Renilla(TU) | Module for the expression of luciferase driven by a GB_SynP promoter with A1 R1, A3 mSlDFR, and an A2 part with three times the target for gRNA1, and the constitutive expression of renilla and P19. |
| GB3331 | Omega1_R1:G1abc.4:mDFR:luc:t35s +P19(TU) +Renilla(TU) | Module for the expression of luciferase driven by a GB_SynP promoter with A1 R1, A3 mSlDFR, and an A2 part with three times the target for gRNA1, and the constitutive expression of renilla and P19. |
| GB3332 | Omega1_R1:G1abc.5:mDFR:luc:t35s +P19(TU) +Renilla(TU) | Module for the expression of luciferase driven by a GB_SynP promoter with A1 R1, A3 mSlDFR, and an A2 part with three times the target for gRNA1, and the constitutive expression of renilla and P19. |
| GB3408 | pUPD2_mSlCHS | Minimal promoter of slCHS1 gene (Solyc09g091510) containing 62bp upstream the transcription start site and the 5'UTR region. |
| GB3409 | pUPD2_mSlANS | Minimal promoter of slANS gene (Solyc10g076660) containing 62bp upstream the transcription start site and the 5'UTR region. |
| GB3410 | pUPD2_mNtTA29 | Minimal promoter of NtTA29 gene containing 62bp upstream the transcription start site and the 5'UTR region (from GB1477 sequence). |
| GB3411 | pUPD2_mAt2S3 | Minimal promoter of At2S3 gene containing 84bp upstream the transcription start site and the 5'UTR region (from GB0029 sequence). |
| GB3412 | pUPD2_mNtPCPS2 | Minimal promoter of NtPCPS2 gene containing 62bp upstream the transcription start site and the 5'UTR region (from GB1027 sequence). |
| GB3413 | pUPD2_mPcPAF | Minimal promoter of PcPAF gene containing 62bp upstream the transcription start site and the 5'UTR region (from FB029 sequence). |
| GB3414 | pUPD2_mAnGPDA | Minimal promoter of AnGPDA gene containing 62bp upstream the transcription start site and the 5'UTR region (from FB007 sequence). |
| GB3520 | Alpha1_R1:G1abc.1:mSlCHS:luc:t35s | TU for the expression of firefly luciferase driven by a GB_SynP promoter with A1 R1 and A3 mSlCS parts, and an A2 part containing three times the gRNA-1DFR (gRNA1) target sequence. |
| GB3521 | Alpha1_R1:G1abc.1:mSlANS:luc:t35s | TU for the expression of luciferase driven by a GB_SynP promoter with A1 R1, A3 mSlANS, and an A2 part containing three times the gRNA-1DFR (gRNA1) target sequence. |
| GB3522 | Alpha1_R1:G1abc.1:mNtTA29:luc:t35s | TU for the expression of luciferase driven by a GB_SynP promoter with A1 R1, A3 mNtTA29, and an A2 part containing three times the gRNA-1DFR (gRNA1) target sequence. |
| GB3523 | Alpha1_R1:G1abc.1:mAt2S3:luc:t35s | TU for the expression of luciferase driven by a GB_SynP promoter with A1 R1, A3 mAt2S3, and an A2 part containing three times the gRNA-1DFR (gRNA1) target sequence. |
| GB3524 | Alpha1_R1:G1abc.1:mNtPCPS2:luc:t35s | TU for the expression of luciferase driven by a GB_SynP promoter with A1 R1, A3 mNtPCPS2, and an A2 part containing three times the gRNA-1DFR (gRNA1) target sequence. |
| GB3525 | Alpha1_R1:G1abc.1:mPcPAF:luc:t35s | TU for the expression of luciferase driven by a GB_SynP promoter with A1 R1, A3 mPcPAF, and an A2 part containing three times the gRNA-1DFR (gRNA1) target sequence. |
| GB3526 | Alpha1_R1:G1abc.1:mAnGPDA:luc:t35s | TU for the expression of luciferase driven by a GB_SynP promoter with A1 R1, A3 mAnGPDA, and an A2 part containing three times the gRNA-1DFR (gRNA1) target sequence. |
| GB3527 | Omega1_R1:G1abc.1:mSlCHS:luc:t35s +P19(TU) +Renilla(TU) | Module for the expression of luciferase driven by a GB_SynP promoter with A1 R1, A3 mSlCHS, and an A2 part with three times the target for gRNA1, and the constitutive expression of renilla and P19. |
| GB3528 | Omega1_R1:G1abc.1:mSlANS:luc:t35s +P19(TU) +Renilla(TU) | Module for the expression of luciferase driven by a GB_SynP promoter with A1 R1, A3 mSlANS, and an A2 part with three times the target for gRNA1, and the constitutive expression of renilla and P19. |
| GB3529 | Omega1_R1:G1abc.1:mNtTA29:luc:t35s +P19(TU) +Renilla(TU) | Module for the expression of luciferase driven by a GB_SynP promoter with A1 R1, A3 mNtTA29, and an A2 part with three times the target for gRNA1, and the constitutive expression of renilla and P19. |

| | | |
|---|---|---|
| GB3530 | Omega1_R1:G1abc.1:mAt2S3:luc:t35s +P19(TU) +Renilla(TU) | Module for the expression of luciferase driven by a GB_SynP promoter with A1 R1, A3 mAt2S3, and an A2 part with three times the target for gRNA1, and the constitutive expression of renilla and P19. |
| GB3531 | Omega1_R1:G1abc.1:mNtPCPS2:luc: t35s +P19(TU) +Renilla(TU) | Module for the expression of luciferase driven by a GB_SynP promoter with A1 R1, A3 mNtPCPS2, and an A2 part with three times the target for gRNA1, and the constitutive expression of renilla and P19. |
| GB3532 | Omega1_R1:G1abc.1:mPcPAF:luc: t35s +P19(TU) +Renilla(TU) | Module for the expression of luciferase driven by a GB_SynP promoter with A1 R1, A3 mPcPAF, and an A2 part with three times the target for gRNA1, and the constitutive expression of renilla and P19. |
| GB3533 | Omega1_R1:G1abc.1:mAnGPDA:luc: t35s +P19(TU) +Renilla(TU) | Module for the expression of luciferase driven by a GB_SynP promoter with A1 R1, A3 mAnGPDA, and an A2 part with three times the target for gRNA1, and the constitutive expression of renilla and P19. |
| GB4435 | Alpha1_R2:G1ab.1:min2S3:luc:t35s | TU for the expression of firefly luciferase driven by a GB_SynP promoter with A1 R2 and A3 mAt2S3 parts, and an A2 part containing the target for the gRNA-1DFR (gRNA1) twice. |
| GB4436 | Omega1_R2:G1ab.1:min2S3:luc:t35s +P19(TU) +Renilla(TU) | Module for the expression of luciferase driven by a GB_SynP promoter with A1 R2, A3 mAt2S3, and an A2 part with the target for gRNA1 twice, and the constitutive expression of renilla and P19. |
| GB4437 | Alpha1_p35s:HispS:t35s | TU for constitutive expression of HispS enzyme |
| GB4438 | Alpha1_p35s:Luz:t35s | TU for constitutive expression of Luz enzyme |
| GB4439 | Alpha2_p35s:H3H:t35s | TU for constitutive expression of H3H enzyme |
| GB4440 | Alpha1_R1:G1abc.1:minPCPS2:HispS:t35s | TU for inducible expression of HispS enzyme, using a GB_SynP promoter with one copy of the target sequence for gRNA1 (1x). |
| GB4441 | Alpha1_R1:G1abc.1:minPCPS2:Luz:t35s | TU for inducible expression of Luz enzyme, using a dCasEV-regulated synthetic promoter with one copy of the target sequence for gRNA1DFR (1x). |
| GB4442 | Alpha2_R1:G1abc.1:minPCPS2:H3H:t35s | TU for inducible expression of H3H enzyme, using a dCasEV-regulated synthetic promoter with one copy of the target sequence for gRNA1DFR (1x). |
| GB4443 | Alpha1_R2:G1ab.1:min2S3:HispS:t35s | TU for inducible expression of HispS enzyme, using a dCasEV-regulated synthetic promoter with two copies of the target sequence for gRNA1DFR (2x). |
| GB4444 | Alpha1_R2:G1ab.1:min2S3:Luz:t35s | TU for inducible expression of Luz enzyme, using a dCasEV-regulated synthetic promoter with two copies of the target sequence for gRNA1DFR (2x). |
| GB4445 | Alpha2_R2:G1ab.1:min2S3:H3H:t35s | TU for inducible expression of H3H enzyme, using a dCasEV-regulated synthetic promoter with two copies of the target sequence for gRNA1DFR (2x). |
| GB4446 | Alpha1_R3:G1aG2b.1:minDFR:HispS:t35s | TU for inducible expression of HispS enzyme, using a dCasEV-regulated synthetic promoter with three copies of the target sequence for gRNA1DFR (3x). |
| GB4447 | Alpha1_R3:G1aG2b.1:minDFR:Luz:t35s | TU for inducible expression of Luz enzyme, using a dCasEV-regulated synthetic promoter with three copies of the target sequence for gRNA1DFR (3x). |
| GB4448 | Alpha2_R3:G1aG2b.1:minDFR:H3H:t35s | TU for inducible expression of H3H enzyme, using a dCasEV-regulated synthetic promoter with three copies of the target sequence for gRNA1DFR (3x). |
| GB4449 | Omega1_35s:Luz:t35s+35s:H3H:t35s | Module for the constitutive expression of Luz and H3H enzymes |
| GB4450 | Omega2_35s:HispS:t35s+35s:CPH:tNOS | Module for the constitutive expression of HispS and CPH enzymes |
| GB4451 | Omega1_1x:Luz+1x:H3H | Module for the inducible expression of (1x)Luz and (1x)H3H enzymes |
| GB4452 | Omega1_1x:Luz+2x:H3H | Module for the inducible expression of (1x)Luz and (2x)H3H enzymes |
| GB4453 | Omega1_1x:Luz+3x:H3H | Module for the inducible expression of (1x)Luz and (3x)H3H enzymes |
| GB4454 | Omega1_2x:Luz+1x:H3H | Module for the inducible expression of (2x)Luz and (1x)H3H enzymes |
| GB4455 | Omega1_2x:Luz+2x:H3H | Module for the inducible expression of (2x)Luz and (2x)H3H enzymes |
| GB4456 | Omega1_2x:Luz+3x:H3H | Module for the inducible expression of (2x)Luz and (3x)H3H enzymes |
| GB4457 | Omega1_3x:Luz+1x:H3H | Module for the inducible expression of (3x)Luz and (1x)H3H enzymes |

| | | |
|---|---|---|
| GB4458 | Omega1_3x:Luz+2x:H3H | Module for the inducible expression of (3x)Luz and (2x)H3H enzymes |
| GB4459 | Omega1_3x:Luz+3x:H3H | Module for the inducible expression of (3x)Luz and (3x)H3H enzymes |
| GB4460 | Omega2_1x:HispS+35s:CPH | Module for the inducible expression of (1x)HispS and the constitutive expression of CPH |
| GB4461 | Omega2_2x:HispS+35s:CPH | Module for the inducible expression of (2x)HispS and the constitutive expression of CPH |
| GB4462 | Omega2_3x:HispS+35s:CPH | Module for the inducible expression of (3x)HispS and the constitutive expression of CPH |
| GB4497 | Alpha2_35s:Luz.t35s+35s:H3H:t35s +35s:HispS:t35s+35s:CPH:tNOS | Module for the constitutive expression of Luz, H3H, HispS and CPH enzymes |
| GB4498 | Alpha2_1x:Luz+1x:H3H+1x:HispS +35sCPH | Module for the inducible expression of (1x)Luz, (1x)H3H and (1x)HispS enzymes, and the constitutive expression of CPH |
| GB4499 | Alpha2_1x:Luz+2x:H3H+1x:HispS +35s:CPH | Module for the inducible expression of (1x)Luz, (2x)H3H and (1x)HispS enzymes, and the constitutive expression of CPH |
| GB4500 | Alpha2_1x:Luz+3x:H3H+1x:HispS +35s:CPH | Module for the inducible expression of (1x)Luz, (3x)H3H and (1x)HispS enzymes, and the constitutive expression of CPH |
| GB4501 | Alpha2_2x:Luz+1x:H3H+1x:HispS +35s:CPH | Module for the inducible expression of (2x)Luz, (1x)H3H and (1x)HispS enzymes, and the constitutive expression of CPH |
| GB4502 | Alpha2_2x:Luz+2x:H3H+1x:HispS +35s:CPH | Module for the inducible expression of (2x)Luz, (2x)H3H and (1x)HispS enzymes, and the constitutive expression of CPH |
| GB4503 | Alpha2_2x:Luz+3x:H3H+1x:HispS +35s:CPH | Module for the inducible expression of (2x)Luz, (3x)H3H and (1x)HispS enzymes, and the constitutive expression of CPH |
| GB4504 | Alpha2_3x:Luz+1x:H3H+1x:HispS +35s:CPH | Module for the inducible expression of (3x)Luz, (1x)H3H and (1x)HispS enzymes, and the constitutive expression of CPH |
| GB4505 | Alpha2_3x:Luz+2x:H3H+1x:HispS +35s:CPH | Module for the inducible expression of (3x)Luz, (2x)H3H and (1x)HispS enzymes, and the constitutive expression of CPH |
| GB4506 | Alpha2_3x:Luz+3x:H3H+1x:HispS +35s:CPH | Module for the inducible expression of (3x)Luz, (3x)H3H and (1x)HispS enzymes, and the constitutive expression of CPH |
| GB4507 | Alpha2_1x:Luz+1x:H3H+2x:HispS +35s:CPH | Module for the inducible expression of (1x)Luz, (1x)H3H and (2x)HispS enzymes, and the constitutive expression of CPH |
| GB4508 | Alpha2_1x:Luz+2x:H3H+2x:HispS +35s:CPH | Module for the inducible expression of (1x)Luz, (2x)H3H and (2x)HispS enzymes, and the constitutive expression of CPH |
| GB4509 | Alpha2_1x:Luz+3x:H3H+2x:HispS +35s:CPH | Module for the inducible expression of (1x)Luz, (3x)H3H and (2x)HispS enzymes, and the constitutive expression of CPH |
| GB4510 | Alpha2_2x:Luz+1x:H3H+2x:HispS +35s:CPH | Module for the inducible expression of (2x)Luz, (1x)H3H and (2x)HispS enzymes, and the constitutive expression of CPH |
| GB4511 | Alpha2_2x:Luz+2x:H3H+2x:HispS +35s:CPH | Module for the inducible expression of (2x)Luz, (2x)H3H and (2x)HispS enzymes, and the constitutive expression of CPH |
| GB4512 | Alpha2_2x:Luz+3x:H3H+2x:HispS +35s:CPH | Module for the inducible expression of (2x)Luz, (3x)H3H and (2x)HispS enzymes, and the constitutive expression of CPH |
| GB4513 | Alpha2_3x:Luz+1x:H3H+2x:HispS +35s:CPH | Module for the inducible expression of (3x)Luz, (1x)H3H and (2x)HispS enzymes, and the constitutive expression of CPH |
| GB4514 | Alpha2_3x:Luz+2x:H3H+2x:HispS +35s:CPH | Module for the inducible expression of (3x)Luz, (2x)H3H and (2x)HispS enzymes, and the constitutive expression of CPH |
| GB4515 | Alpha2_3x:Luz+3x:H3H+2x:HispS +35s:CPH | Module for the inducible expression of (3x)Luz, (3x)H3H and (2x)HispS enzymes, and the constitutive expression of CPH |
| GB4516 | Alpha2_1x:Luz+1x:H3H+3x:HispS +35s:CPH | Module for the inducible expression of (1x)Luz, (1x)H3H and (3x)HispS enzymes, and the constitutive expression of CPH |
| GB4517 | Alpha2_1x:Luz+2x:H3H+3x:HispS +35s:CPH | Module for the inducible expression of (1x)Luz, (2x)H3H and (3x)HispS enzymes, and the constitutive expression of CPH |
| GB4518 | Alpha2_1x:Luz+3x:H3H+3x:HispS +35s:CPH | Module for the inducible expression of (1x)Luz, (3x)H3H and (3x)HispS enzymes, and the constitutive expression of CPH |
| GB4519 | Alpha2_2x:Luz+1x:H3H+3x:HispS +35s:CPH | Module for the inducible expression of (2x)Luz, (1x)H3H and (3x)HispS enzymes, and the constitutive expression of CPH |
| GB4520 | Alpha2_2x:Luz+2x:H3H+3x:HispS +35s:CPH | Module for the inducible expression of (2x)Luz, (2x)H3H and (3x)HispS enzymes, and the constitutive expression of CPH |
| GB4521 | Alpha2_2x:Luz+3x:H3H+3x:HispS +35s:CPH | Module for the inducible expression of (2x)Luz, (3x)H3H and (3x)HispS enzymes, and the constitutive expression of CPH |
| GB4522 | Alpha2_3x:Luz+1x:H3H+3x:HispS +35s:CPH | Module for the inducible expression of (3x)Luz, (1x)H3H and (3x)HispS enzymes, and the constitutive expression of CPH |

| | | |
|---|---|---|
| GB4523 | Alpha2_3x:Luz+2x:H3H+3x:HispS +35s:CPH | Module for the inducible expression of (3x)Luz, (2x)H3H and (3x)HispS enzymes, and the constitutive expression of CPH |
| GB4524 | Alpha2_3x:Luz+3x:H3H+3x:HispS +35s:CPH | Module for the inducible expression of (3x)Luz, (3x)H3H and (3x)HispS enzymes, and the constitutive expression of CPH |
| GB4558 | Omega2_eGFP+P19+35s:Luz+ 35s:H3H:t35s+35s:HispS:t35s+ 35s:CPH:tNOS | Module for the constitutive expression of Luz, H3H, HispS, CPH eGFP and P19 proteins |
| GB4559 | Omega2_eGFP+P19+1x:Luz+1x:H3H +1x:HispS+35s:CPH | Module for the inducible expression of (1x)Luz, (1x)H3H and (1x)HispS enzymes, and the constitutive expression of CPH, eGFP and P19 |
| GB4560 | Omega2_eGFP+P19+1x:Luz+2x:H3H +1x:HispS+35s:CPH | Module for the inducible expression of (1x)Luz, (2x)H3H and (1x)HispS enzymes, and the constitutive expression of CPH, eGFP and P19 |
| GB4561 | Omega2_eGFP+P19+1x:Luz+3x:H3H +1x:HispS+35s:CPH | Module for the inducible expression of (1x)Luz, (3x)H3H and (1x)HispS enzymes, and the constitutive expression of CPH, eGFP and P19 |
| GB4562 | Omega2_eGFP+P19+2x:Luz+1x:H3H +1x:HispS+35s:CPH | Module for the inducible expression of (2x)Luz, (1x)H3H and (1x)HispS enzymes, and the constitutive expression of CPH, eGFP and P19 |
| GB4563 | Omega2_eGFP+P19+2x:Luz+2x:H3H +1x:HispS+35s:CPH | Module for the inducible expression of (2x)Luz, (2x)H3H and (1x)HispS enzymes, and the constitutive expression of CPH, eGFP and P19 |
| GB4564 | Omega2_eGFP+P19+2x:Luz+3x:H3H +1x:HispS+35s:CPH | Module for the inducible expression of (2x)Luz, (3x)H3H and (1x)HispS enzymes, and the constitutive expression of CPH, eGFP and P19 |
| GB4565 | Omega2_eGFP+P19+3x:Luz+1x:H3H +1x:HispS+35s:CPH | Module for the inducible expression of (3x)Luz, (1x)H3H and (1x)HispS enzymes, and the constitutive expression of CPH, eGFP and P19 |
| GB4566 | Omega2_eGFP+P19+3x:Luz+2x:H3H +1x:HispS+35s:CPH | Module for the inducible expression of (3x)Luz, (2x)H3H and (1x)HispS enzymes, and the constitutive expression of CPH, eGFP and P19 |
| GB4567 | Omega2_eGFP+P19+3x:Luz+3x:H3H +1x:HispS+35s:CPH | Module for the inducible expression of (3x)Luz, (3x)H3H and (1x)HispS enzymes, and the constitutive expression of CPH, eGFP and P19 |
| GB4568 | Omega2_eGFP+P19+1x:Luz+1x:H3H +2x:HispS+35s:CPH | Module for the inducible expression of (1x)Luz, (1x)H3H and (2x)HispS enzymes, and the constitutive expression of CPH, eGFP and P19 |
| GB4569 | Omega2_eGFP+P19+1x:Luz+2x:H3H +2x:HispS+35s:CPH | Module for the inducible expression of (1x)Luz, (2x)H3H and (2x)HispS enzymes, and the constitutive expression of CPH, eGFP and P19 |
| GB4570 | Omega2_eGFP+P19+1x:Luz+3x:H3H +2x:HispS+35s:CPH | Module for the inducible expression of (1x)Luz, (3x)H3H and (2x)HispS enzymes, and the constitutive expression of CPH, eGFP and P19 |
| GB4571 | Omega2_eGFP+P19+2x:Luz+1x:H3H +2x:HispS+35s:CPH | Module for the inducible expression of (2x)Luz, (1x)H3H and (2x)HispS enzymes, and the constitutive expression of CPH, eGFP and P19 |
| GB4572 | Omega2_eGFP+P19+2x:Luz+2x:H3H +2x:HispS+35s:CPH | Module for the inducible expression of (2x)Luz, (2x)H3H and (2x)HispS enzymes, and the constitutive expression of CPH, eGFP and P19 |
| GB4573 | Omega2_eGFP+P19+2x:Luz+3x:H3H +2x:HispS+35s:CPH | Module for the inducible expression of (2x)Luz, (3x)H3H and (2x)HispS enzymes, and the constitutive expression of CPH, eGFP and P19 |
| GB4574 | Omega2_eGFP+P19+3x:Luz+1x:H3H +2x:HispS+35s:CPH | Module for the inducible expression of (3x)Luz, (1x)H3H and (2x)HispS enzymes, and the constitutive expression of CPH, eGFP and P19 |
| GB4575 | Omega2_eGFP+P19+3x:Luz+2x:H3H +2x:HispS+35s:CPH | Module for the inducible expression of (3x)Luz, (2x)H3H and (2x)HispS enzymes, and the constitutive expression of CPH, eGFP and P19 |
| GB4576 | Omega2_eGFP+P19+3x:Luz+3x:H3H +2x:HispS+35s:CPH | Module for the inducible expression of (3x)Luz, (3x)H3H and (2x)HispS enzymes, and the constitutive expression of CPH, eGFP and P19 |
| GB4577 | Omega2_eGFP+P19+1x:Luz+1x:H3H +3x:HispS+35s:CPH | Module for the inducible expression of (1x)Luz, (1x)H3H and (3x)HispS enzymes, and the constitutive expression of CPH, eGFP and P19 |
| GB4578 | Omega2_eGFP+P19+1x:Luz+2x:H3H +3x:HispS+35s:CPH | Module for the inducible expression of (1x)Luz, (2x)H3H and (3x)HispS enzymes, and the constitutive expression of CPH, eGFP and P19 |
| GB4579 | Omega2_eGFP+P19+1x:Luz+3x:H3H +3x:HispS+35s:CPH | Module for the inducible expression of (1x)Luz, (3x)H3H and (3x)HispS enzymes, and the constitutive expression of CPH, eGFP and P19 |
| GB4580 | Omega2_eGFP+P19+2x:Luz+1x:H3H +3x:HispS+35s:CPH | Module for the inducible expression of (2x)Luz, (1x)H3H and (3x)HispS enzymes, and the constitutive expression of CPH, eGFP and P19 |
| GB4581 | Omega2_eGFP+P19+2x:Luz+2x:H3H +3x:HispS+35s:CPH | Module for the inducible expression of (2x)Luz, (2x)H3H and (3x)HispS enzymes, and the constitutive expression of CPH, eGFP and P19 |
| GB4582 | Omega2_eGFP+P19+2x:Luz+3x:H3H +3x:HispS+35s:CPH | Module for the inducible expression of (2x)Luz, (3x)H3H and (3x)HispS enzymes, and the constitutive expression of CPH, eGFP and P19 |
| GB4583 | Omega2_eGFP+P19+3x:Luz+1x:H3H +3x:HispS+35s:CPH | Module for the inducible expression of (3x)Luz, (1x)H3H and (3x)HispS enzymes, and the constitutive expression of CPH, eGFP and P19 |
| GB4584 | Omega2_eGFP+P19+3x:Luz+2x:H3H +3x:HispS+35s:CPH | Module for the inducible expression of (3x)Luz, (2x)H3H and (3x)HispS enzymes, and the constitutive expression of CPH, eGFP and P19 |
| GB4585 | Omega2_eGFP+P19+3x:Luz+3x:H3H +3x:HispS+35s:CPH | Module for the inducible expression of (3x)Luz, (3x)H3H and (3x)HispS enzymes, and the constitutive expression of CPH, eGFP and P19 |

**A**

```
G1ab.5        --CCATCTTCTCTCACCAACCAGTCatcgaattcgtccgtgttcatgttatatatgcaca   58
G1ab.4        --CCATCTTCTCTCACCAACCAGTCaagaccagggggctcgccgcgttggctaatcctg   58
G1ab.1        -tCCCTCTTCTCTCACCAACCAGTCagtcgtgaaagtcatagtaccctgggtaccaactt   59
G1abc.4       -tCCCTCTTCTCTCACCAACCAGTCacgaccgcttcacgctaaggtgctggccacgtgct   59
G1abc.3       -tCCCTCTTCTCTCACCAACCAGTCagttgggttcacccgggtcggacctgagtcgacca   59
G1abc.1       -tCCCTCTTCTCTCACCAACCAGTCgtatatccgtttcaattcgtttttctcgtctacag   59
G1abc.2       -tCCCTCTTCTCTCACCAACCAGTCaacgccccgtaccgatgctgaacaagtcgatgcag   59
G1abc.5       -tCCCTCTTCTCTCACCAACCAGTCcttactgtatgagtagtaatttgctcgagatgtcg   59
G1a.4         --GCTGTATCTAATAGAATCTTCGGgcgtacatcctacctgaggtctgtggccccgtggt   58
G1a.6         --GCTGTATCTAATAGAATCTTGGGgctagcttaggcatcattgttacccatagcgcggc   58
G1ab.3        taCCATCTTCTCTCACCAACCAGTCtctgaattcgcagactcagtaagacacggtctagc   60
G1a.5         ---CTGTATCTAATAGAATCTTGGGtgaattcatattactgtcataaccgctcagttcgt   57
G1a.2         --GCTGTATCTAATAGAATCTTAGGgagaattcagatctacctcctaaggcactacgaag   58
G1ab.2        --CCATCTTCTCTCACCAACCAGTCagtgagctcctgtgctcggtggtcatttggtatgg   58
G1a.1         --GCTGTATCTAATAGAATCTTGGGacatgggcgtttggcactaccgacacgaacctcag   58
G1a.3         -aGCTGTATCTAATAGAATCTTAGGacatgggcgtttggcactaccgacacgaacctcag   59
                 *    ***   *  * *

G1ab.5        agCCGTCTTCTCTCACCAACCAGTCaacaggctaggatataatgctgaagcccttcccca   118
G1ab.4        gtCCCTCTTCTCTCACCAACCAGtcacatcttgtaatgaatattcagtagaaaatttgtg   118
G1ab.1        acCCGTCTTCTCTCACCAACCAGTCcgctggttcctgggtgagctcgagactcgtggtga   119
G1abc.4       aaCCTTCTTCTCTCACCAACCAGTCtggcgttgatgtggatcgactctatagCCATCTTC   119
G1abc.3       agCCTTCTTCTCTCACCAACCAGTCgcgaccgcagaacttacgctcgagggtCCGTCTTC   119
G1abc.1       ccCCTTCTTCTCTCACCAACCAGTCacctgcactaccttactgcggtccgcaCCGTCTTC   119
G1abc.2       gcCCTTCTTCTCTCACCAACCAGTCcaatcccgaggcctgacccgacatatcCCATCTTC   119
G1abc.5       gtCCTTCTTCTCTCACCAACCAGTCtaacgcgtcgtatctacgtcacgacgaCCGTCTTC   119
G1a.4         caCCTTCTTCTCTCACCAACCAGTCggagcgtaatcagccgtatccagcaacactacgct   118
G1a.6         gtCCTTCTTCTCTCACCAACCAGTCgcgacggtcctgacctgctactcttccgcaagaat   118
G1ab.3        tgCCTTCTTCTCTCACCAACCAGTCggagtccggtgtagcgaaagatcaaggcgaccta   120
G1a.5         cgCCCTCTTCTCTCACCAACCAGTCgattcctgcgtaataacaatctgtagccacgcaa-   116
G1a.2         gaCCATCTTCTCTCACCAACCAGTCcacgggctcgacatccaggctgggattttgacatg   118
G1ab.2        tcCCCTCTTCTCTCACCAACCAGTCtctatcactggaaatcggacgtgaggtaggat---   115
G1a.1         ttCCTTCTTCTCTCACCAACCAGTCagaaacactggaaatgggacgtgaggtaggat---   115
G1a.3         ttCCGTCTTCTCTCACCAACCAGTCctcgccagagctcgtcagcatactcgaagaat---   116
                ** ********************

G1ab.5        agcgttcagggtgggatttgctacaacttccga-------      151
G1ab.4        ttagaaggacgagtcaccatgtaccaaaagcga-------      151
G1ab.1        cagctcttcatacatagagcggccgcgtcgaac-------      152
G1abc.4       TCTCACCAACCAGTCcgaggccgttcgtt-----------      148
G1abc.3       TCTCACCAACCAGTCtccccggttatctc-----------      148
G1abc.1       TCTCACCAACCAGTCccagggaggacctc-----------      148
G1abc.2       TCTCACCAACCAGTCgtagctaactatgt-----------      148
G1abc.5       TCTCACCAACCAGTCagggctagaattac-----------      148
G1a.4         a----tctggtcatatcataagattccgcgagctcaa---      151
G1a.6         g-----gtggtccagccgtccaactcagctcaacatag--      151
G1ab.3        ggtagcaaccgccggcttcggcggtaag-----ggaat--      153
G1a.5         ----gacttcggcgtccttgggtggggacgctatgaat--      150
G1a.2         gagaggctggtaattgttttggtgtggctgaat-------      151
G1ab.2        ----ggcttgtcctttcattcgttgccgactgagctctta      151
G1a.1         ----cggttgtcctatctttcgttgccgactgcgcagaat      151
G1a.3         ----caaggcaggtcaattcgcactgtgagagtcgaagtg      152
```

**B**

```
G1d.1         ------------------------------------------------------------   0
G1a.7         ------------------------------------------------------------   0
G3aG2b.1      ------------------------------------------------------------   0
G1b.1         ------------------------------------------------------------   0
G1e.1         CCGTCTTCTCTCACCAACCAGTCatacctggagctgtaccgttattgcgctgcatagatg   60

G1d.1         ------------------------------------------------------------   0
G1a.7         ------------------------------------------------------------   0
G3aG2b.1      ------------------------------------------------------------   0
G1b.1         ---------------------------------------------------CATCTTCTC   9
G1e.1         cagtgctgctcttatcacatttgtttcgacgacagccgccttcgcagtttcctcagacac   120

G1d.1         -----------------------------------------taGCTGTATCTAATAGAAT   19
G1a.7         -------------GCTGTATCTAATAGAATCTTCGGctattagtggctgcggcaaaatat   47
G3aG2b.1      -------------GCTGTATCTAATAGAATCTTCGGctattagtggctgcggcaaaatat   47
G1b.1         TCACCAACCAGTCcttattgtaggcagaggcacgccctattagtggctgcggcaaaatat   69
G1e.1         ttaagaataagcgcttattgtaggcagaggcacgccctattagtggctgcggcaaaatat   180
                                                             *     * *  **

G1d.1         CTTCGGagcgaattcctgtgctcggtggtcaaatggtatcgtgCCGTCTTCTCTCACCAA   79
G1a.7         cttctaagcgaatCCGTCTTCTCTCACCAACCAGTCtcctgtgctcggtggtcaaatggt   107
G3aG2b.1      cttctaagcgaatCCAGAAACCCGCGGCTCAGTGGCtcctgtgctcggtggtcaaatggt   107
G1b.1         c-----------------------ttctaagcgaattcctgtgctcggtggtcaaatggt   106
G1e.1         c-----------------------ttctaagcgaattcctgtgctcggtggtcaaatggt   217
              *                                   *   ****    *  **  *

G1d.1         CCAGTCgctgcgtaatcagccgtatccagcaacactacgctatc  123
G1a.7         atcgtggctgcgtaatcagccgtatccagcaacactacgctatc  151
G3aG2b.1      atcgtggctgcgtaatcagccgtatccagcaacactacgctatc  151
G1b.1         atcgtggctgcgtaatcagccgtatccagcaacactacgctatc  150
G1e.1         atcgtggctgcgtaatcagccgtatccagcaacactacgctatc  261
                 ** **************************************
```

**Figure S1. Alignment of A2 parts sequences included in the GB_SynP collection**. (A) Alignment of the three A2 parts series including different repetitions of the gRNA1 target (G1a.N, G1ab,N and G1abc.N). (B) Alignment of the A2 part series including the target for gRNA1 at different positions (G1d.1, G1a.7, G1b.1 and G1e.1) and the G3a.1 part that contains the same sequence as G1a.7 but replacing the gRNA1 target by gRNA3 target. Alignments were performed using CLUSTAL-Omega (v1.2.4). Capital letters denote the target sequences for the different gRNAs.

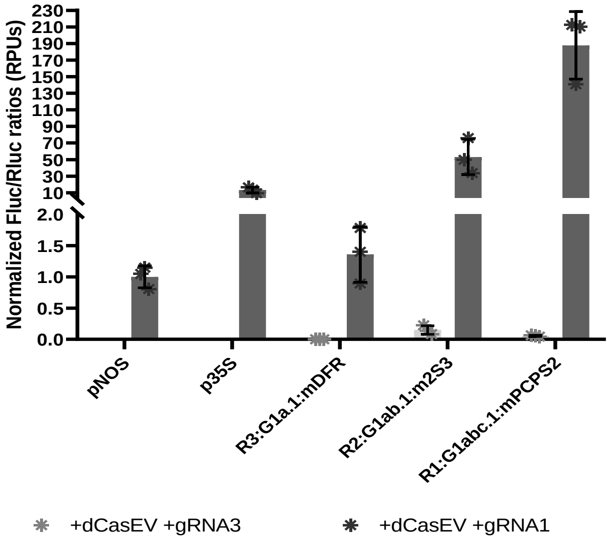


**Figure S2. Expression range of the GB_SynP promoters used for regulation of LUZ pathway.** Normalized (Fluc/Rluc) expression levels of Nicotiana benthamiana leaves transiently expressing a luciferase reporter gene (Fluc) under the regulation of promoter R3:G1a.1:mDFR (1x gRNA-target), R2:G1ab.1:m2S3 (2x gRNA-target) or R1:G1abc.1:mPCPS2 (3x gRNA-target). Luciferase under *NOS* and *CaMV35s* promoters were included as references. Letters denote statistically significance between (activated) promoters (t-test, alpha=0.05). Error bars represent the average values ± SD (n=3).

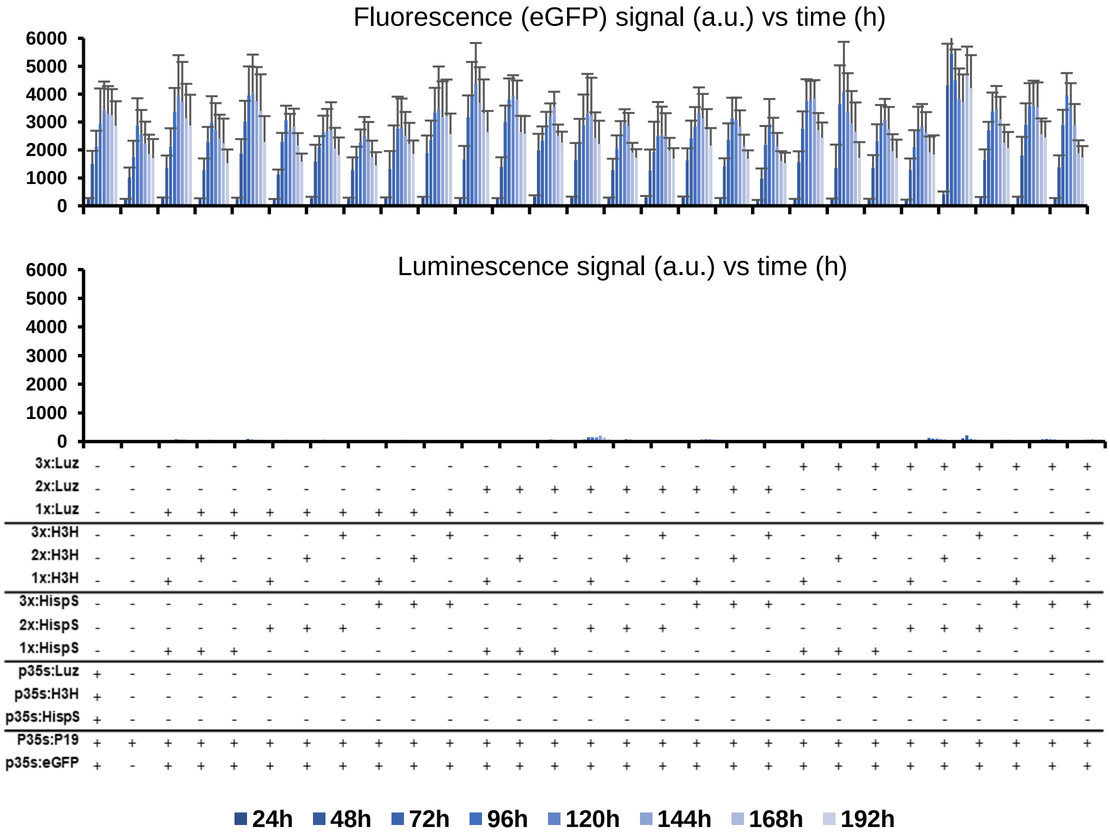


**Figure S3. Basal expression of the Time-course experiment including the 27 constructs expressing the LUZ pathway transiently in *Nicotiana benthamiana* leaves under the regulation of 1x, 2x or 3x GB_SynP promoters.** Fluorescence and luminescence signals corresponds to the co-infiltration in *Nicotiana benthamiana* leaves of each LUZ pathway construct with dCasEV2.1 and an irrelevant gRNA (gRNA3). Error bars represent the average values ± SD (n=12).
